# New genomic resources for an invasive mite pest expand the known diversity of pesticide resistance mechanisms and guide development of KASP diagnostic markers

**DOI:** 10.1101/2025.11.17.688773

**Authors:** Joshua A. Thia, Kelly Richardson, Courtney Brown, Aston Arthur, Svetlana Micic, Alan Lord, Neil D. Young, Sunita B. Sumanam, Jia Chang, Bill C.H. Chang, Shujun Wei, Robin B. Gasser, Thomas L. Schmidt, Paul A. Umina, Ary A. Hoffmann

## Abstract

The redlegged earth mite, *Halotydeus destructor*, is an invasive pest of grain and pasture crops in Australia. Australian populations are increasingly developing resistance to broad-spectrum pesticides. Here, we assembled a chromosome-level genome for *H. destructor*, comprising nine chromosomes and totalling approximately 46 Mb. We also generated a genomic dataset from 190 populations collected across thousands of kilometres of its invasive Australian range and across four years (2018, 2021, 2022, and 2023), and used this dataset to develop KASP (Kompetitive allele-specific PCR) markers for identifying resistance mutations. We combined pooled whole-genome sequencing with resistance testing for organophosphate pesticides, chlorpyrifos and omethoate, and the pyrethroid pesticide bifenthrin.

Target-site resistance to chlorpyrifos was strongly associated with the common *ace* G119S mutation. A second mutation, *ace* A201S, was also common, although its association with chlorpyrifos or omethoate resistance was less clear. Target-site resistance to bifenthrin was strongly associated with the common para L1024F mutation. We also detected several rare target-site mutations that likely contribute to resistance in specific populations, including F331Y in *ace*, and M918T, M918V, L925I, F1020S, and F1538I in *para*. Copy number variation in *ace* showed mixed associations with resistance, potentially reflecting the co-occurrence of variable gene copies and target-site mutations segregating within and among populations. Finally, we observed putative compositional and structural differences in genomic regions containing carboxylesterases with high *ace*-like homology, a group of genes that might be involved in organophosphate metabolism.

This study advances the genomic and molecular resources available for *H. destructor*, and represents the most comprehensive investigation to date of resistance mechanisms and their spatiotemporal distribution in this invasive pest. Our data showed that pesticide resistance in *H. destructor* likely involves many genes. Nevertheless, target-site mutations are likely to remain key markers for rapid resistance diagnostics, for which our KASP markers will provide valuable tools.

## INTRODUCTION

Genomic approaches have greatly advanced many aspects of pest management. For example, genomic studies have illuminated invasion pathways, dispersal capacity, spatial scales of population structure, and admixture events that promote invasiveness (Schmidt et al., 2023; Tepolt & Palumbi, 2020; Zhang et al., 2020). They have also helped identify the genetic architecture underlying adaptation of pests to the local environment (Anderson et al., 2018; Kreiner et al., 2022; Zhang et al., 2023). Such insights are beneficial to pest management because they provide a clearer picture of the demographic and evolutionary processes operating in pest populations, and they help identify functionally important genetic variation in pest organisms (Anderson et al., 2018; Tepolt et al., 2021). This knowledge can inform strategies to limit the spread of pests, their pathogens, and economically harmful traits, such as pesticide resistance (Anderson et al., 2018; Cao et al., 2025; Kreiner et al., 2022). Additionally, genomic data can be used to design genetic monitoring tools for different ecotypes, strains, or resistance mutations (Cao et al., 2025; Thia et al., 2025). Curating reference genomes and spatiotemporally sampling populations for genomic analyses is therefore key to establishing a rich set of genetic resources that can inform pest management strategies (Fritz, 2022; Thia, Hoffmann, et al., 2021).

The evolution of pesticide resistance poses a major challenge in pest management through the loss of chemical control options. In many cases, pesticide resistance is outpacing the registration of new chemical products (Sparks & Nauen, 2015; Umina, McDonald, et al., 2019). At the same time, the range of approved pesticide options is narrowing due to concerns about environmental impacts (FAO, 2022). A key part in managing pesticide resistance is through resistance monitoring activities. The goals of monitoring will depend on the resistance context, and include pre-emptively identifying early signs of resistance, quantifying the extent to which resistance has spread, and tracking changes in resistance following the implementation of management strategies (Bird, 2015; Cao et al., 2025).

Molecular diagnostic markers provide an important tool for resistance monitoring by reducing costs associated with phenotypic screening. One emerging technology for pesticide resistance screening is the Kompetitive allele-specific PCR (KASP) assay system (Shen et al., 2023; Shi et al., 2023). KASP is a fluorescence-based, single-plex genotyping assay that uses allele-specific forward primers with unique tails and a common reverse primer to selectively amplify target Single Nucleotide Polymorphisms (SNPs) (Semagn et al., 2014). During PCR, competitive binding of the allele-specific primers leads to the incorporation of fluor-labelled oligonucleotides, enabling post-PCR discrimination of genotypes through endpoint fluorescence measurement (Semagn et al., 2014). KASP was originally developed for plant breeding programs (Semagn et al., 2014) but has also been applied to pesticide resistance monitoring in arthropods to screen candidate resistance alleles. An additional advantage of KASP is its flexibility for use with pooled DNA samples (Brusa et al., 2023; Shi et al., 2023). This approach not only enables estimation of population-level allele frequencies, but is also particularly valuable for arthropods, where small body sizes can make individual DNA extractions challenging.

In this study, we expand the genomic resources available for the redlegged earth mite, *Halotydeus destructor* (Tucker, 1925) (Trombidiformes: Penthaleidae), and use them to develop new resistance markers. *Halotydeus destructor* is an invasive pest in Australia, first introduced around 1917, after which it rapidly spread across temperate southern regions during the 1920s (Ridsdill-Smith, 1997). It is a major pest of grain and pasture crops and can cause yield losses of up to 80% (Ridsdill-Smith, 1997). For decades, control has relied primarily on pesticides, particularly organophosphates and pyrethroids applied as spray treatments (Ridsdill-Smith, 1997), and neonicotinoids as seed treatments (Umina, Arthur, et al., 2019).

To date, organophosphate and pyrethroid resistance has been reported in Australian *H. destructor*. Resistance first emerged in the early 2000s and is now present across large areas of Australia’s west and parts of its eastern regions (Arthur et al., 2021; Umina, 2007; Umina & Hoffmann, 1999). Resistance can spread via long-distance dispersal but has also been shown locally at a farm-scale (Thia, Korhonen, et al., 2023; Yang et al., 2020). However, previous studies (Hill et al., 2016; Thia, Korhonen, et al., 2023; Weeks et al., 1995; Yang et al., 2020) have been insufficiently powered to fully characterise the distribution of genetic variation at different spatial scales.

A range of mechanisms may underpin resistance in *H. destructor*. Target-site mutations in the protein targets of pesticides appear important. For organophosphates, three mutations, G119S, A201S, and F331Y, in the acetylcholinesterase gene (*ace*), are candidate polymorphisms segregating in field populations of *H. destructor* (Thia, Korhonen, et al., 2023; Thia, Umina, et al., 2023). However, previous studies have not fully evaluated the relative importance of these mutations due to limitations in sampling design (Thia, Korhonen, et al., 2023; Thia, Umina, et al., 2023). Metabolic mechanisms are also likely to be important in organophosphate resistance. Current hypotheses include amplification of the *ace* gene itself (increasing target protein product) as well as expansion of a group of carboxylesterases that exhibit high *ace*-like homology (Thia, Korhonen, et al., 2023; Thia, Umina, et al., 2023). In contrast, for pyrethroids, a single mutation, L1024F, in the *para*-like voltage-gated sodium channel gene (*para*), has a strong association with resistance (Edwards et al., 2018), although metabolic mechanisms might also contribute (Yang et al., 2020).

Our study had four major goals. The first goal was to develop a new chromosome-level assembly for *H. destructor*, as an improvement on the existing scaffold-level draft assembly (Thia, Korhonen, et al., 2023). The second goal was to develop a large population genomic dataset that could provide better resolution of genetic variation and pesticide resistance polymorphisms in *H. destructor* at the broad landscape level (thousands of kilometres). The third goal was to develop a set of KASP markers from candidate target-site mutations for monitoring resistance. The fourth goal was to explore the diversity of *ace*-like carboxylesterases within and among populations using long-read data generated from select field populations.

## METHODS

### Sample collection

Population samples for resistance screening were conducted between 2018 to 2023 as part of a broad monitoring program across southern Australia in both western and eastern regions. The numbers of sample populations were *n* = 6 in 2018, *n* = 47 in 2021, *n* = 92 in 2022, and *n* = 45 in 2023 (*n* = 190 in total). The distribution of sampled populations is illustrated in Figure 1; note, two populations from WA could not be plotted due to lack of geographic coordinate data. Mites were collected by suction using a Stihl SH55 blower vacuum (Stihl, Waiblingen, Germany) with a fine gauze mesh inserted at the end of the vacuum tube. Once collected, mites were placed into plastic containers with vegetation and paper-towelling, and transported back to the laboratory, where they were placed in a fridge at 4°C until phenotypic testing.

**Figure 1.**
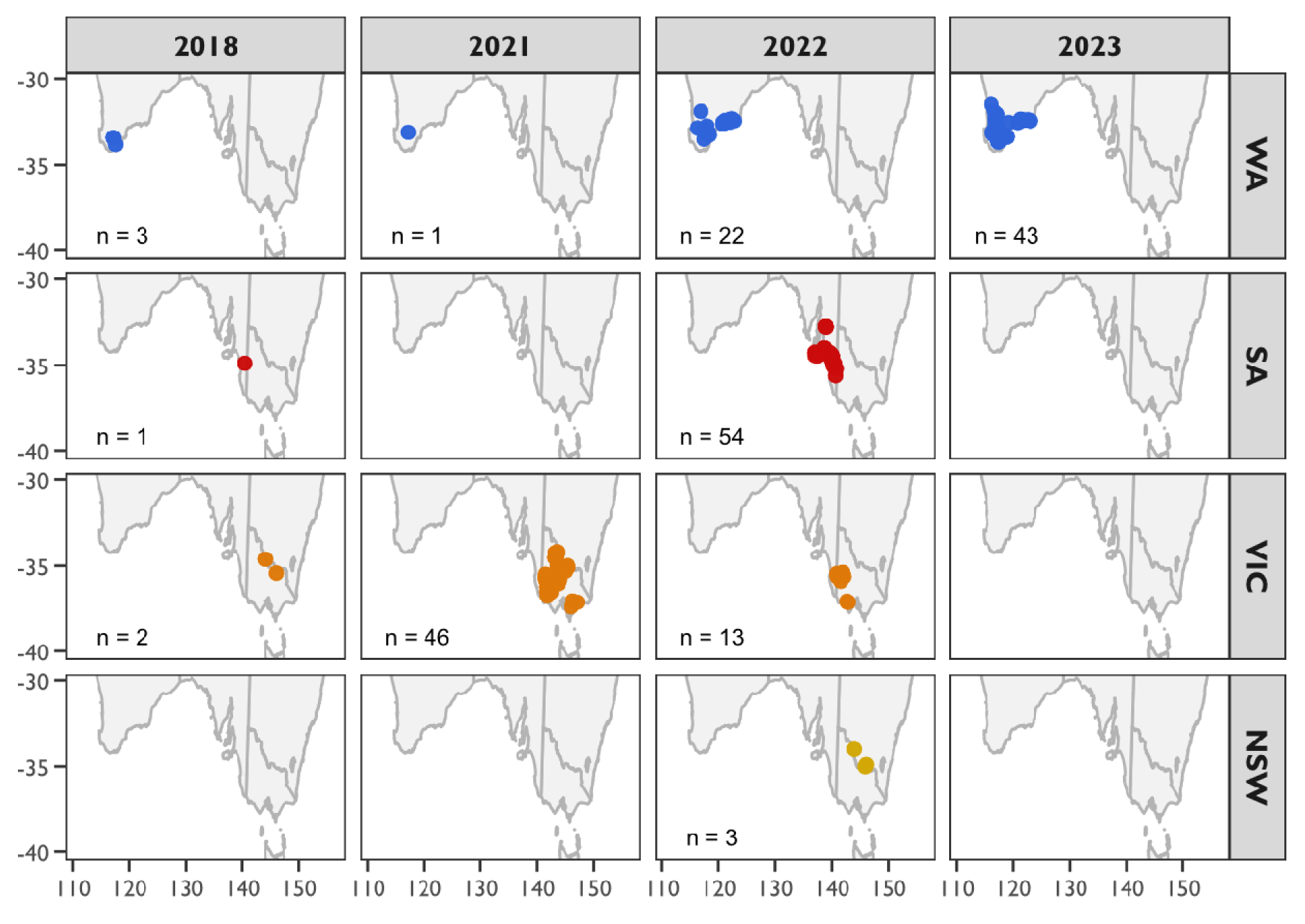
Spatiotemporal sampling of *Halotydeus destructor* for this study. Points represent populations sampled. Panels separate populations by Australian state (rows) and collection years (columns), with number of sampled population (*n*) printed in the bottom left corner. Note, two populations collected from 2023 in WA are not plotted due to unavailable geographic coordinate data (top right corner panel: plotted populations is *n* = 43, but total analysed number is *n* = 45).

Phenotypic resistance data for each population was obtained by screening mites from each sampled population alongside a known pesticide-susceptible population against discriminatory doses of three commonly used pesticides: the organophosphates chlorpyrifos and omethoate, and the pyrethroid bifenthrin. Screening used discriminating doses of 0.021 g/L (chlorpyrifos), 0.0058 g/L (omethoate), and 0.1 g/L (bifenthrin), as identified in previous work (Arthur et al., 2021). Phenotypic screening was undertaken by pouring the pesticide solution of each chemical into a 15 mL plastic vial and swirling to ensure complete coverage, discarding excess solution. Six vials per chemical were coated and then left to dry upside down overnight on a rack. Control vials were treated in the same manner using water. To aid the spread of the pesticide and ensure even coating within the plastic vials, the surfactant, Tween, was added at 0.01% into the water used to generate pesticide dilutions. Once vials were dry, approximately eight *H. destructor* were placed in each vial, along with two to three leaves of common vetch (*Vicia sativa* L), which provided humidity and food. Vials were then sealed with a screw cap lid and place at 18°C for 24 hours. Mites were then scored as alive (moving freely), incapacitated (inhibited movement) or dead (no movement over a 5 second period) (following Umina, 2007). Mites used for generating population genomic data (see section ‘Pooled whole-genome sequencing’) were independent of those used for resistance phenotyping as above; these represent two random samples of mites taken from the same population.

An older collection of *H. destructor* was used for refining the original scaffold-level draft genome into a chromosome-level assembly (see section ‘Hi-C sequencing’). This collection was made in 2021 from the Colbinabbin area in Victoria (−36.5901, 144.7994), at a site known to harbour a population of pesticide-susceptible mites (Thia et al., 2022). Mites were collected from the field as described above, returned to the laboratory for identification and sorting, and then stored in 100% ethanol at −20°C.

### Pooled whole-genome sequencing

Genomic DNA was extracted from pooled samples for all 190 *H. destructor* populations with phenotypic data. We aimed to pool 60 mites per population, although our final dataset comprised pools with a range of 13 to 60 mites and a mean of 57; 83% of pools comprised 60 mites. Mites were homogenised using a TissueLyser III (Qiagen) with three 2 mm glass beads and extracted with either the NEB Monarch® Genomic DNA kit or the Qiagen EZ2 DNA Tissue kit; in both cases, we followed the manufacturer protocols except that we extended the lysis time to be overnight (∼18 hours). Pooled DNA samples were sent to Novogene (Novogene Co. Ltd., Singapore) for library preparation and Illumina short-read sequencing. We aimed for 1.5 Gb of sequence data per sample because the *H. destructor* genome is ∼50 Mb, equivalent to a sequencing depth of 30×. We obtained a mean of 13,676,053 raw reads per library, with a range of 9,550,221 to 24,070,912 raw reads.

### Hi-C sequencing

Mites from a 2021 collection from Colbinabbin, Victoria, were used to generate Hi-C data. A total of 1,200 mites of mixed sex were pooled into a single sample. The pooled mites were processed using the Arima High Coverage Hi-C kit (L/N2309050008; part no. A160162 v01; Arima Genomics, Carlsbad, CA, USA) according to manufacturer’s protocols. The final library was quality and integrity checked using the TapeStation 4200 system Genomic DNA ScreenTape assay (Agilent, CA, USA) and paired-end sequenced on a NovaSeq X instrument at the Australian Genome Research Facility (AGRF Ltd., Peter MacCallum Cancer Centre facility, Melbourne, Australia). The short-reads were stored in the FASTQ format for quality check and subsequent analysis.

### Chromosome-level assembly

Our Hi-C data was used to complete the publicly available draft genome (NCBI Assembly ASM2275052v1, GenBank GCA_022750525.1, BioProject PRJNA756307) to a chromosome-level assembly. This original draft genome was assembled with mites sampled from a population in Wantirna South (Melbourne, Victoria) (Thia, Korhonen, et al., 2023). This population is genetically very similar to the Colbinabbin population used to generate the Hi-C data (*F*_ST_ = 0.02) (Thia, Korhonen, et al., 2023).

The raw Illumina short-reads were trimmed with *fastp* v0.23.4 (Chen et al., 2018) to remove residual adapter sequence, trim to a mean quality of 20, and retain only reads with length ≥60 bp. This produced 528,135,528 trimmed and paired reads. Trimmed reads were mapped to the draft genome using *chromap* v0.3.1 (H. Zhang et al., 2021) and filtered with *samtools* v1.16 (Danecek et al., 2021; Li et al., 2009) to retain reads with mapping scores >30. The Hi-C assembly with mapped reads was performed using *yahs* v1.2.2 (Zhou et al., 2023) and *juicer pre* v1.9.9 (Durand, Shamim, et al., 2016). The preliminary assembly was visualised and manually curated with *juicebox* v2.17.00 (Durand, Robinson, et al., 2016). The curated assembly was then finalised using *juicer post* v1.9.9 (Durand, Shamim, et al., 2016). Gaps in the assembly were filled with *dentist* v4.0.0 (Durand, Robinson, et al., 2016; Ludwig et al., 2022) using PacBio long-reads generated from a previous study (Thia, Korhonen, et al., 2023) (NCBI Sequence Read Archive SRR15698716).

Custom R code was applied to ‘clean’ the assembly; large gaps of ‘N’ were standardised to a constant size, chromosomes and scaffolds ordered by size and renamed. Repetitive elements were identified using *red* v 2018.09.10 (Girgis, 2015), isolated with *bedtools* v2.30.0 (Danecek et al., 2021), and annotated with *repeatmodeler* v2.0.4 (Flynn et al., 2020). The completeness of the final genome was assessed using *busco* v5.7.1 (Simão et al., 2015) with the Arthropoda ortholog database (‘genome’ mode), and redundancy was assessed with a self-to-self alignment using *minimap2* v2.26 (Li, 2016) and visualised with the *pafr* v0.0.2 R package (Winter, 2020).

Gene annotation of our chromosome-level genome was achieved using Illumina short-read transcriptome data from a previous study (Thia, Korhonen, et al., 2023) (NCBI SRA SRR20755148 and SRR20755147). Reads were trimmed with *fastp* (same as Hi-C reads). Trimmed reads were mapped using *hisat2* v2.2.1 (Kim et al., 2019) and filtered with *samtools* to retain alignments with mapping score >30. Gene models were predicted using *braker* v3.0.6 (Gabriel et al., 2024). Gene annotation file formats were manipulated with *gffread* 0.12.7 (Pertea & Pertea, 2020) and *agat* v1.0.0 (Dainat, 2022). A combination of *biopython* v1.79 (Chapman & Chang, 2000; Cock et al., 2009), *augustus* v3.5.0 (Stanke et al., 2006), and *seqkit* v2.3.1 (W. Shen et al., 2016) was applied to extract the longest protein-coding transcript for each gene. A combination of *blastp* from the *blast+* suit v2.13.0 (Camacho et al., 2009) and *eggnog* v2.1.12 (Cantalapiedra et al., 2021) was used to assign functions to annotated genes.

### Comparative genomics

We performed a comparative analysis between our new *H. destructor* chromosome-level assembly and that of the two-spotted spider mite, *Tetranychus urticae* (Koch, 1836) (Trombidiformes: Tetranychidae). We chose *T. urticae* because it is the most well studied mite in the order Trombidiformes and belongs to one of the most recently derived families, the Tetranychidae (Klimov et al., 2018; Qin, 1996; Szudarek-Trepto et al., 2022). In contrast, *H. destructor* is a member of one of the most ancestral families, the Penthaleidae (Klimov et al., 2018; Qin, 1996; Szudarek-Trepto et al., 2022).

We used the *T. urticae* NCBI GenBank reference genome, GCA_039701765.1, in our comparative analysis. This genome has been assembled to the chromosome level (the RefSeq *T. urticae* genome is only at the scaffold level). Assembly completeness was assessed using *busco* with the Arthropoda ortholog database (‘genome’ mode). Synteny between *H. destructor* and *T. urticae* was performed through a combination of *minimap2* to align the chromosome assemblies, with visualisations performed in R using *pafr* and *circlize* (Gu et al., 2014). The two genomes were quite divergent from each other, and syntenic blocks were small and (or) did not have high similarity. We visualised aligned regions that were ≥1,000 bp with ≥70% identity. The final circle plot was edited with *Inkscape* (Inkscape Project, 2025).

### Population genomics

#### Bioinformatic processing

Pooled whole-genome sequencing reads were trimmed using *fastp* to remove residual adapter sequence, attain a mean quality of Q20, and retain only reads with length ≥100 bp. This retained a mean of 13,419,672 trimmed reads per pool, with a range of 9,367,035 to 23,656,783 reads. Trimmed reads were mapped with *bowtie2* v2.4.5 (Langmead & Salzberg, 2012) and *samtools* was used to retain alignments with mapping score >20 and to calculate coverage in 1,000 bp windows. Variants were called using *bcftools mpileup* v1.15.1 with a combination of *bcftools filter* and *vcftools* v0.1.16 (Danecek et al., 2011) applied to perform an initial filtering of variants to retain SNPs with a minimum depth of 20 reads, and with missing data ≤5%. These variants were imported in R using *genomalicious* v0.7.11 (Thia, 2024) and filtered further to retain those with a minor allele frequency of 5% across populations. This resulted in 804,510 SNPs, of which 772,909 were biallelic and 31,601 were triallelic. Functional annotation of SNP variants was performed with *snpeff* v5.1f (Cingolani et al., 2012). All downstream analyses were performed in R.

#### Genetic diversity

All post-filtered SNPs were used to calculate genomic heterozygosity, π, as the estimator π (Ferretti et al., 2013), using a simplified calculation that we derived previously (Thia, Korhonen, et al., 2023):

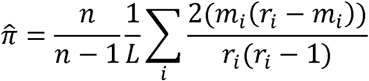

Here, *n* is the number of haploid individuals in a pool, *L* is the length of a focal genomic window, *i* is the index of a focal nucleotide position nested within a genomic window, *r_i_* is the read depth for position *i*, and *m* is the observed alternate read count (ranging from 1 to *r_i_* – 1). SNP variants and genomic coverage were filtered separately for each pool. SNPs and genomic windows were only retained if they had a minimum depth of 20 reads. The size of the genomic window (≤1000 bp) was used to represent *L*. We calculated a genome-wide 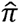 for each population by taking the median across genomic windows.

#### Population genetic structure

For computational efficiency, we reduced our SNP dataset to biallelic SNPs spaced at 5 kb intervals across the genome. We quantified the magnitude of genetic differentiation among all populations, and between pairs of populations, using Weir & Cockerham’s *F*_ST_ (1984). Expected SNP heterozygosity for biallelic loci was calculated as *H*_E_ = 1 – *H*_O_, where *H*_O_ = *p*^2^ + *q*^2^. The package *genomalicious* was used calculate the variance components, *a*, *b* and *c* (Weir & Cockerham, 1984), to estimate *F*_ST_.

We tested for patterns of isolation-by-distance by correlating geographic distance (kilometers) with genetic differentiation (*F*_ST_). We excluded 2018 populations from this analysis due to a limited sample size. We analysed populations sampled in 2021, 2022, and 2023 separately to avoid confounding spatial and temporal genetic differentiation.

Qualitative patterns of population genetic structure were visualised using principal components analysis (PCA), conducted on all populations across all years and focusing on the two main PC axes that accounted for most of the variation (Supplementary Figure S1). The R function *prcomp* was used to perform the PCA on allelic covariances (arguments: ‘scale=FALSE, center=TRUE’). The screeplot of explained variances was considered to infer the number of significant axes of population genetic structure (Patterson et al., 2006; Thia, 2023).

#### Target-site mutations

We took a subset of our SNP dataset consisting of those loci found in the target resistance genes, *ace* (for organophosphates) and *para* (for pyrethroids) and the *nAChR-*β*1* subunit (for neonicotinoids). We used a dataset without minor allele frequency (MAF) filtering to capture mutations that might be segregating at a low frequency. For target-site mutation analysis, we allowed for the inclusion of triallelic SNPs by calculating a bipartite MAF that compared reference alleles vs the sum of all alternate alleles. Herein, we used canonical (reference) species codon numbering to make our work directly comparable to previous literature: *Torpedo californica* for *ace*, *Muscae domestica* for *para*, and *Myzus persicae* for *nAChR*.

We examined genotype-phenotype associations for candidate organophosphate and pyrethroid resistance mutations as potential KASP markers. We used linear regression to quantify associations for three mutations, *ace* G119S, *ace* A201S, and *para* L1024. We selected these known target-site mutations as the major candidates (see Results). Linear regressions were fit using the *lm* function in R, with a Gaussian error distribution, and assessed the proportion of mites surviving pesticide exposure as predicted by the mutant allele frequency.

#### Copy number variation in *ace*

Another putative mechanism involved in organophosphate resistance is increased copy number of the *ace* gene. We sampled 4 non-overlapping 1,000 bp windows that spanned the canonical *ace* in our *H. destructor* assembly. As a proxy for *ace* copy number, we calculated the standardised read depth for each population by dividing the read depth in these windows by the genomic average, and took the mean. We tested the hypothesis that survival following pesticide exposure was associated with copy number by fitting the model:

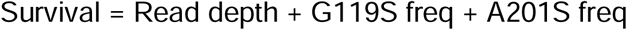

Here, the response ‘Survival’ is the proportion of mites surviving pesticide exposure. The continuous fixed effect predictors are ‘Read depth’, as the proxy for copy number, ‘G119S freq’ as the frequency of the G119S mutation, and ‘A201S freq’ as the frequency of the A201S mutation. All predictors were normalised to a mean of 0 and variance of 1. Models were fit using R’s *glm* function with a quasibinomial error distribution with equal weights across observations. Significance was assessed using the *Anova* function from the *car* package using Type II sums-of-squares. This allowed us to assess the contribution of each predictor after accounting for the other terms in the model, which would reveal how much variation in survival was explained by read depth after accounting for the frequencies of known target-site mutations. We fit models separately for each organophosphate pesticide (chlorpyrifos and omethoate) and state (WA, SA, and VIC), excluding NSW because of small sample size.

For quality control, we also explored whether carboxylesterase genes with homology to the canonical *ace* gene might influence estimates of copy number (see also Methods Section ‘Expansion of ace-like carboxylesterase genes’ below). For this assessment, we used the population (RLEM202) with the highest observed standardised depth at the canonical *ace* gene (2.57-fold relative to genomic background). We extracted the reads mapping to the canonical *ace* gene using *samtools*, then attempted to map these reads to the sequences of carboxylesterase genes that were annotated with *ace*-like homologies using *bowtie2*. If *ace*-like genes do not interfere with copy number estimates, no cross-mapping should be observed.

### KASP marker development

Based on our population genomic analyses, we identified the G119S mutation in *ace*, and the L1204F mutation in *para*, as candidates for KASP marker development. KASP genotyping assays were performed using reagents from LGC Biosearch Technologies (Middlesex, United Kingdom) with support from GeneWorks Pty Ltd (South Australia, Australia). The assay ID numbers were KBS-2300-00-436054101 (*ace* G119S mutation) and KBS-2300-00-436067771 and KBS-2300-00-436067772 (*para* L1024F mutations, Type I and 2, respectively). We tested the ability of these KASP primer sets to detect the presence of the target-site mutations and recover the allele frequencies observed in our population genomic data. As positive controls, we designed a series of synthetic DNA molecules, gBlocks, in *geneious prime* (Geneious, 2021). These gBlocks were synthesized by Integrated DNA Technologies (Coralville, Iowa, USA).

KASP assays were performed on the same pooled DNA extractions used to generate the whole-genome sequencing data. A subset of 90 pools across four Australian states were selected to capture a range of resistant and susceptible populations with a broad geographic spread. DNA samples were diluted to a range of 0.2−2 ng/μL. We diluted gBlocks to be at a final working concentration of 0.05 ng/μL. We found that gBlocks amplified effectively at this concentration, and this low concentration helped reduce assay costs. PCR free water provided a negative control. Each reaction comprised 2.5 μL sample, 2.5 μL 2X KASP-TF Master Mix, and 0.07 μL KASP primer mix. We ran the KASP assays on a Roche LightCycler® 480 system (Roche Diagnostics Pty. Ltd). The cycling conditions involved four stages: (1) 94°C for 15 minutes; (2) 10 cycles of 94°C for 20 seconds, followed by 60 seconds starting at 68°C that dropped 0.6°C each cycle, ending at 62°C by the 10^th^ cycle; (3) 26 cycles of 94°C for 20 secs, followed by 55°C for 60 seconds; and (4) 37°C for 1 minute.

Fluorescence data was imported into R for analysis (association testing) and visualisation. We assessed the correlation between HEX signals and pool-seq allele frequencies to determine the ability of the KASP assays to quantify allele frequencies in pooled DNA samples. We also fitted a logistic regression for each mutation to estimate the slopes of these relationships (*b*_1_). Models were fit using *glm* function in R, with a quasi-binomial error distribution, with equal weights across observations, and using the allele frequencies as the response variable and the HEX signal as a continuous fixed effect.

### Expansion of *ace*-like carboxylesterase genes

In addition to copy number variation in the *ace* gene itself, we previously hypothesised that expansion and copy number of carboxylesterases with *ace*-like homology might also contribute to pesticide resistance in *H. destructor* (Thia, Umina, et al., 2023). We explored variation in this group of genes by using long-read data generated from pooled DNA samples from two field collected populations: RLEM102 (susceptible) and RLEM120 (omethoate and bifenthrin resistant). Nanopore libraries were prepared using the NBD 114.24 kit (Oxford Nanopore Technologies) and were sequenced on a Promethion 2 Solo platform by Oz Omics (Melbourne, Australia: https://www.ozomics.com/).

Trimming of the Nanopore long reads was performed using *fastp*, filtering reads to a minimum length of 10,000 bp. We then performed a targeted assembly by mapping reads to our reference genome using *minimap2*. Mapped reads were retained and assembled with *flye2*. Completeness was assessed using *busco* and the Arthropoda dataset. Transcriptomic data from a previous study (Thia, Korhonen, et al., 2023) was mapped to the contigs using *hisat2*, and gene models were called with *braker3*. Functional annotations were performed using a combination of *eggnog* and *blastp*.

Genes with *ace*-like annotations were imported into R for further analysis. These were combined with the canonical *ace* and *ace*-like genes from the reference assembly. All protein sequences were filtered to retain only those with ≥400 amino acid residues. An alignment was performed using the *msa* package (Bodenhofer et al., 2015), which was then filtered to produce a matrix of informative sites (no gaps or invariant sites) with the *genomalicious* package. Amino acid differences were calculated using the *ape* package (Paradis et al., 2004), and a UPGMA tree was produced using the *phangorn* package (Schliep, 2011). The gene tree was rooted on the reference *ace* gene and visualised with the *ggtree* package (Yu et al., 2017).

Our protein alignment revealed some *ace*-like genes within populations that exhibited high similarity (few, or no, amino acid differences at our informative sites). We herein refer to these as ‘clusters’ of similar *ace*-like genes, those with ≤5 amino acid differences. For each cluster, we checked whether they occurred on the same assembled chromosome (for the reference assembly) or contig (for populations RLEM102 and RLEM120). We also examined the annotations around these *ace*-like genes to gain a better understanding of their genomic context—this was also useful for evaluating the presence of split contigs during the assembly.

## RESULTS

### Genome assembly

We assembled a chromosome-level reference genome for *H. destructor* using Hi-C sequence data. This new assembly comprises nine chromosomes (46 Mb) that range from 4.18 Mb to 5.74 Mb, and 57 contigs (3.00 Mb) that range from 1,000 bp to 220,082 bp (Figure 2a). The N50 and N90 were 4.98 and 4.18 Mb, respectively, and the L50 and L90 were five and nine sequences, respectively. This assembly showed low redundancy, as evidenced by a self-to-self alignment (Figure 2b). Annotation of this new assembly identified 13,726 protein coding genes. BUSCO scores for the Arthropoda single-copy ortholog set indicated high completeness: 90.5% of genes were captured by our assembly (of these, 89.6% were single-copy and 0.9% were duplicated); 2.3% of genes were fragmented; 7.2% of genes were missing. A summary of annotated repetitive elements can be found in Appendix 1. Our new assembly and annotations have been submitted to NCBI.

**Figure 2.**
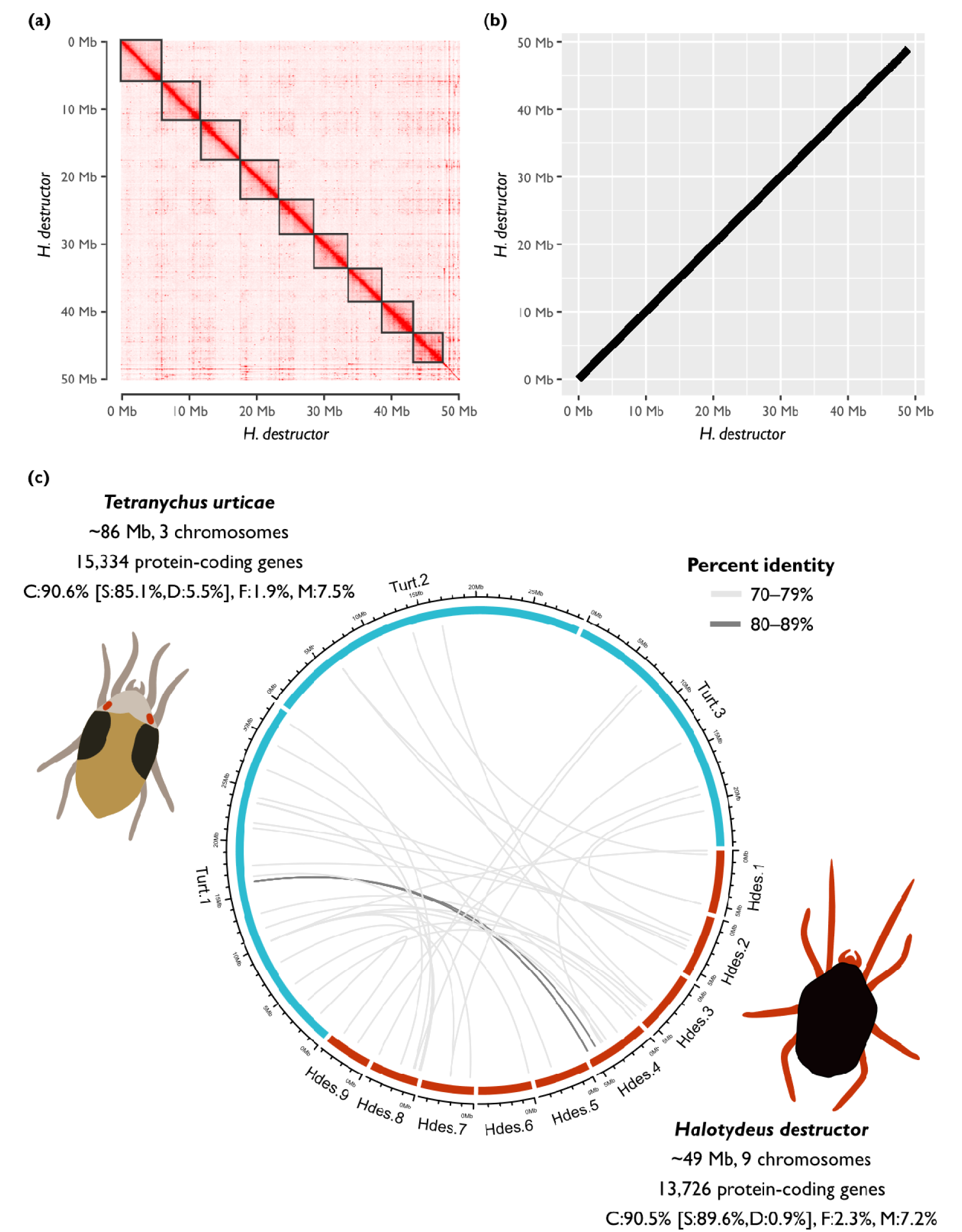
*Halotydeus destructor* chromosome-level assembly and comparative analysis with *Tetranychus urticae*. (a) Hi-C contact heatmap scaffolding the *H. destructor* genome into nine chromosomes. Interaction frequency is shown as a red gradient, with higher frequencies along the diagonal. Boxes delineate chromosome boundaries. (b) Dotplot of the self–self alignment of the *H. destructor* assembly (chromosomes only). The continuous diagonal indicates consistent chromosome alignments with no evidence of discordant matches, suggesting the assembly is not redundant. (c) Comparative analysis of synteny between our *H. destructor* assembly and a *T. urticae* assembly. Chromosome sequences are positioned on the edge of the ring: the nine *H. destructor* chromosomes are labelled as ‘Hdes’ and coloured red, and the three *T. urticae* chromosomes are labelled as ‘Turt’ and coloured blue. The grey ribbon indicates syntenic region between species that are ≥1,000 bp long, with shades of grey indicating percent similarity (see legend). Summary statistics for the assembly of each species are reported at the edges of the ring, including BUSCO percentages for the arthropod database: ‘C’ are complete genes, broken into ‘S’ true single-copies and ‘D’ duplicated copies, ‘F’ fragmented genes, and ‘M’ missing genes.

### Comparative genomics

Annotation of the *T. urticae* assembly (NCBI GenBank GCA_039701765.1) returned 14,991 genes. BUSCO scores for the Arthropoda single-copy ortholog set indicated reasonable completeness: 90.6% of genes were captured by our assembly (of these, 85.4% were single-copy and 5.2% were duplicated); 1.9% of genes were fragmented; 7.5% of genes were missing. These BUSCO scores were comparable to those in our *H. destructor* assembly. There was very little synteny between the *H. destructor* and *T. urticae* genomes (Figure 2c). In total, there were 37 syntenic regions that were ≥1,000 bp and ≥70% similarity. Most of these were small, with a mean syntenic region size of 2,205 bp, and a range of 1,001 bp to 11,797 bp.

### Population genomics

#### Genetic diversity

The genomic heterozygosity, 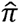, exhibited a median of 0.0073 across the 190 *H. destructor* populations, with a range of 0.0062 to 0.0075 (Figure 3b). Populations from the state of WA had the greatest diversity (median 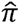 of 0.0074), followed by those from SA (0.0073), and then VIC (0.0072). We sampled only three populations from NSW (0.0073), so we were unable to characterise the distribution of values within this state, nor relate it to other states. A Kruskal-Wallis test among WA, SA, and VIC populations indicated significant differences in 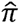 (χ^2^ = 24.7, df = 2, *P* < 0.001).

**Figure 3.**
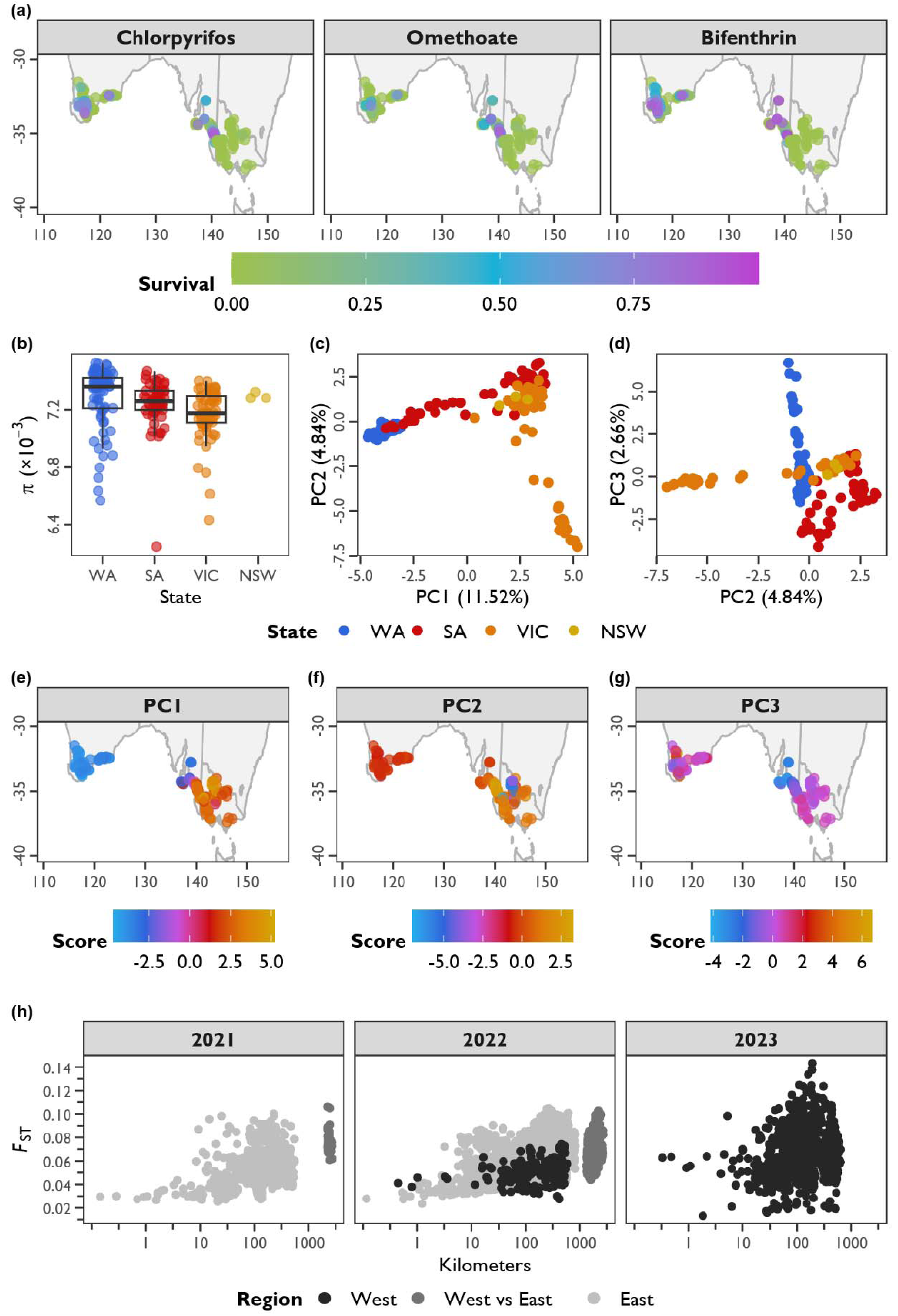
Distribution of resistance and population genetic structure in Australian populations of *Halotydeus destructor*. (a) Maps of resistances in southern Australia. Panels separate different pesticide compounds. Each point is a mite population, coloured by their survival following exposure to discriminating concentrations of each pesticide (see legend). (b) Genomic heterozygosity (π) of populations from each Australian state. States are arranged on the *x*-axis, with genomic heterozygosity on the *y*-axis (note the scale is ×10^−3^). Points represent mite populations, and boxplots summarise the distribution (there is no boxplot for the state of NSW because only three populations were sampled). (c, d) Principle components (PCs) of the nuclear genetic variation, PC1 vs PC2 (c), and PC2 vs PC3 (d). The *x*- and *y*-axes represent the different PC scores. Percentages in the axis labels indicate the amount of allelic variation explained by each PC axis. Points represent mite populations, coloured by state (see legend). (e, f, g) Maps of PC scores for PC1 (e), PC2 (f), and PC3 (g). Points represent mite populations, coloured by their respective PC score (see legend). (h) Isolation-by-distance between population pairs. Geographic distance is on the *x*-axis, measured in kilometers (note: real values have been overlaid onto the log10-transformed scale). Genetic differentiation is on the *y*-axis, measured as *F*_ST_. Points represent distances between population pairs, coloured by region (west or east) (see legend). Panels separate population pairs from different years.

#### Population genetic structure

PCA revealed regional structuring of genetic variation (Figure 3c to 3g). The first two PC axes were the most informative of population structure, based on the ‘elbowing’ of eigenvalues (explained variances) at PC3 (Supplementary Figure S1) (Patterson et al., 2006; Thia, 2023). PC1, PC2, and PC3 explained 11.52%, 4.84%, and 2.66% of the total variation in allele frequencies, respectively. PC1 broadly captured a dimension of genetic variation that exhibited a west-to-east pattern (Figure 3c and 3e). At the ends of the PC1 spectrum were WA populations in the west, and VIC and NSW populations in the east. Populations in SA exhibited a gradient of scores that transitioned from being ‘west-like’ to more ‘east-like’. PC2 captured a dimension of genetic variation describing the variability among SA, VIC, and NSW populations (Figure 3c and 3f). PC3 largely captured a dimension of genetic variation describing variability among WA and SA populations (Figure 3d and 3g).

There was weak to moderate genetic structure among populations. The mean pairwise *F*_ST_ was 0.069, with a range of 0.013 to 0.14 across population pairs. Isolation-by-distance was logarithmic: *F*_ST_ rose very quickly over tens of kilometers between population pairs, and populations separated by hundreds to thousands of kilometers could be similarly differentiated (Figure 3h; see also Supplementary Figure S2). The correlations (*r*) between *F*_ST_ and log_10_-transformed kilometers across all comparisons was *r* = 0.52, 0.42, and 0.12 for the 2021, 2022, and 2023 collections, respectively.

#### Target-site mutations

Screening of non-synonymous mutations in target genes revealed 12 mutations in *ace* (organophosphates), 19 in *para* (pyrethroids), and 1 in *nAChR-*β*1*. In *ace*, 3 mutations are known target-site mutations: G119S, A201S, and F331Y (*T. californica* numbering), but here only G119S and A201S had MAF > 0.05 (Supplementary Table S1). In *para*, 6 mutations are known target-site mutations: M198V, M918T, L925I, F1020Y, L1024F, and F1538I (*M. domestica* numbering), but only L1024F had MAF > 0.05 (Supplementary Table S1). The mutation in *nAChR-*β*1* is not known to contribute to target-site resistance and had MAF < 0.05 (Supplementary Table S1).

Resistance association analysis focused on the *ace* G119S and A201S mutations and the *para* L1024F mutation, given their appreciable MAF and known contributions to resistance in other arthropods (Figure 4). However, all resistance associations for mutations with MAF > 0.05 are reported in Supplementary Figures S3, S4 and S5. We also present population counts of known target-site mutations in Table 1, and their mean allele frequencies in Supplementary Table S1.

**Figure 4.**
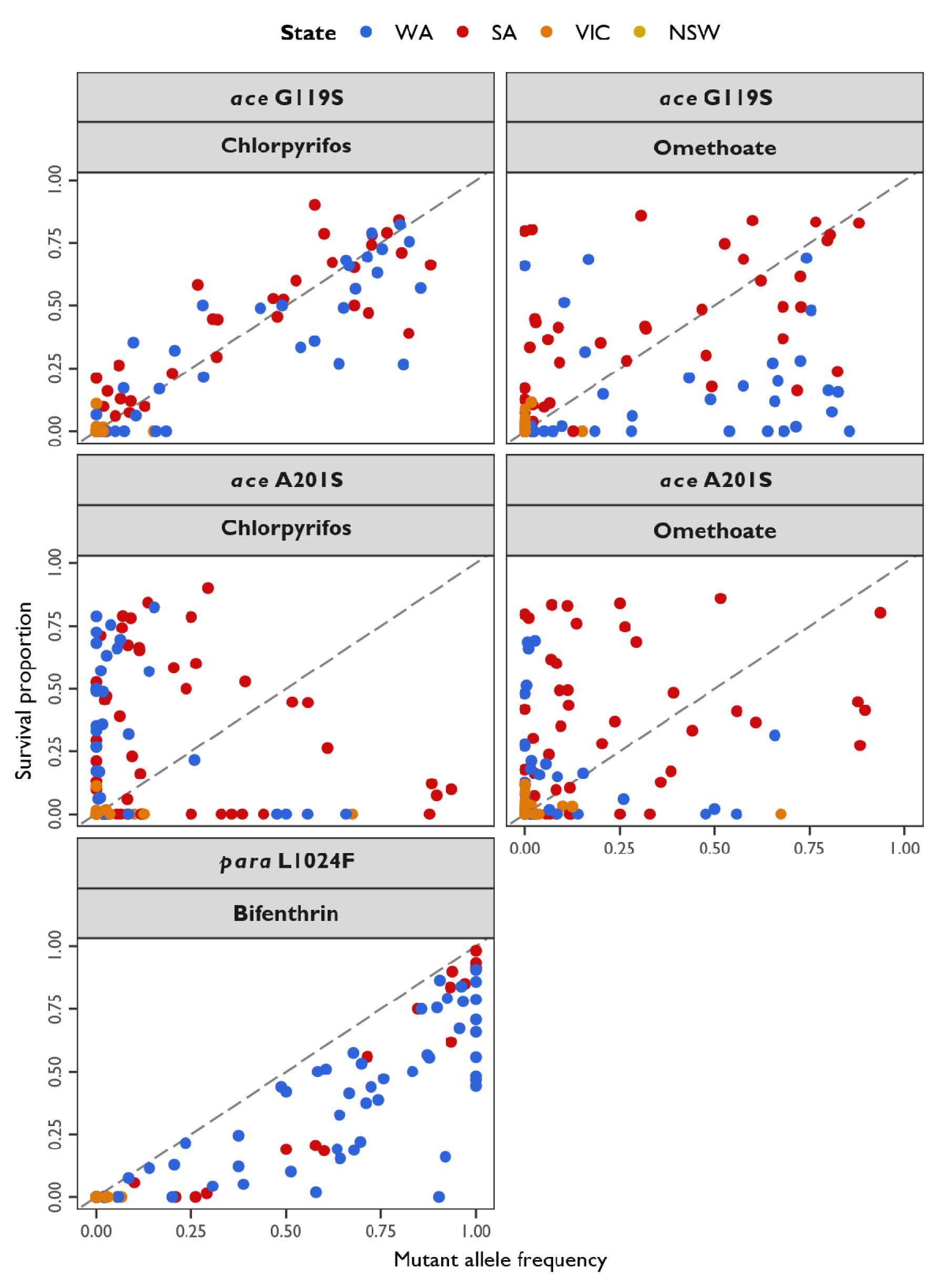
Associations between candidate target-site resistance mutations (G119S, A201S, and L1024F) and population-level resistance in *Halotydeus destructor*. The *x*-axis represents the allele frequency of each mutation, and the *y*-axis represents the proportional survival of mites following exposure to a discriminating concentration of chlorpyrifos, omethoate and bifenthrin. Points represent mite populations, coloured by Australian state (see legend). The dashed line indicates an expected slope of 1. Panels display results for each combination of mutation and pesticide.

**Table 1.** Known target-site mutations in *ace* and *para* genes of *Halotydeus destructor* that occur below a minor allele frequency of 0.05 across all 190 populations sampled. In this table, the number of populations for which the mutations are segregating is reported, along with the mean, minimum and maximum frequency of the mutations across the populations in which they are found. See also Supplementary Table S1 for Australia-wide statistics.

| Gene | Mutation | State | Num.<br>pops. | Mean<br>freq. | Min<br>freq. | Max<br>freq. |
| --- | --- | --- | --- | --- | --- | --- |
| ace | F331Y | WA | 11 | 0.33 | 0.01 | 0.91 |
|  |  | SA | 10 | 0.03 | 0.01 | 0.10 |
|  |  | VIC | 3 | 0.02 | 0.02 | 0.02 |
|  |  | NSW | 1 | 0.03 | 0.03 | 0.03 |
| para | M918T | WA | 18 | 0.07 | 0.01 | 0.20 |
|  |  | SA | 10 | 0.08 | 0.02 | 0.17 |
|  |  | VIC | 3 | 0.04 | 0.02 | 0.07 |
|  |  | NSW | 1 | 0.02 | 0.02 | 0.02 |
|  | L925I | WA | 19 | 0.10 | 0.01 | 0.30 |
|  |  | SA | 4 | 0.03 | 0.02 | 0.04 |
|  |  | VIC | 3 | 0.02 | 0.02 | 0.02 |
|  |  | NSW | 1 | 0.02 | 0.02 | 0.02 |
|  | F1020Y | WA | 10 | 0.11 | 0.02 | 0.31 |
|  |  | SA | 5 | 0.03 | 0.02 | 0.04 |
|  |  | VIC | 2 | 0.02 | 0.02 | 0.02 |
|  |  | NSW | 1 | 0.02 | 0.02 | 0.02 |
|  | F1538I | WA | 8 | 0.10 | 0.03 | 0.41 |
|  |  | SA | 3 | 0.03 | 0.02 | 0.05 |
|  |  | VIC | 1 | 0.03 | 0.03 | 0.03 |

For *ace*, the G119S mutation had a mean frequency of 0.14 (range of 0 to 0.88), and the A201S mutation had a mean frequency of 0.07 (range of 0 to 0.93) (Figure 4: top row). The G119S mutations exhibited a strong association with chlorpyrifos (*b*_0_ = 0.01, *b*_1_ = 0.89, *R*^2^ = 0.87, *p* < 0.001), and a moderate association with the second organophosphate tested, omethoate (*b*_0_ = 0.05, *b*_1_ = 0.50, *R*^2^ = 0.34, *p* < 0.001). The A201S mutation had no association with chlorpyrifos (*b*_0_ = 0.12, *b*_1_ = 0.16, *R*^2^ = 0.01, *p* = 0.101) and a weak association with omethoate (*b*_0_ = 0.08, *b*_1_ = 0.50, *R*^2^ = 0.16, *p* < 0.001) (Figure 4: middle row). The frequency of the G119S mutation was either completely uncorrelated with the frequency of A201S, or negatively correlated with it, depending on the location of the mite population (Supplementary Figure S5). This suggests that these mutations are competing in occurrence and/or are on different haplotypes.

For the *para* L1024F mutation, the mean frequency was 0.24 (range of 0 to 1). This mutation is underpinned by two different alleles: the Type 1 allele is a TTT codon with a mean frequency of 0.2 (range of 0 to 1), and the Type 2 allele is a TTC codon with a mean frequency of 0.04 (range of 0 to 0.76). The association between bifenthrin resistance and the L1024F mutation was strong (*b*_0_ = 0.012, *b*_1_ = 0.68, *R*^2^ = 0.84, *p* < 0.001) (Figure 4: bottom row). All three alleles could segregate within populations. Based on allelic counts, the wild-type TTG codon exhibited a correlation of *r* = −0.84 and −0.52 with the TTT and TTC codons, respectively, and the mutant TTT codon exhibited a correlation of *r* = 0.30 with the mutant TTC codon.

#### Copy number variation in *ace*

Read depth was used as a proxy for *ace* copy number, measured as an average across 1,000 bp genomic windows encompassing the gene (Figure 5a). Copy number variation was substantial among WA and SA populations, but not among VIC populations (Figure 5b). We found mixed support for the hypothesis that *ace* copy number is positively associated with pesticide resistance in WA and SA, after accounting for the frequency of target-site mutations (Supplementary Table S2 & S3). For chlorpyrifos, ‘Read depth’ was a non-significant predictor in WA (χ^2^ = 0.4, df = 1, *P* = 0.524), negatively associated with resistance in SA (χ^2^ = 5.8, df = 1, *P* = 0.016, β = −0.37), and a non-significant predictor in VIC (χ^2^ = 0.02, df = 1, *P* = 0.88). For omethoate, ‘Read depth’ was positively associated with resistance in WA (χ^2^ = 71.8, df = 1, *P* < 0.001, β = 1.27), but was a non-significant predictor in SA (χ^2^ = 0.1, df = 1, *P* = 0.794) and VIC (χ^2^ = 0.2, df = 1, *P* = 0.639).

**Figure 5.**
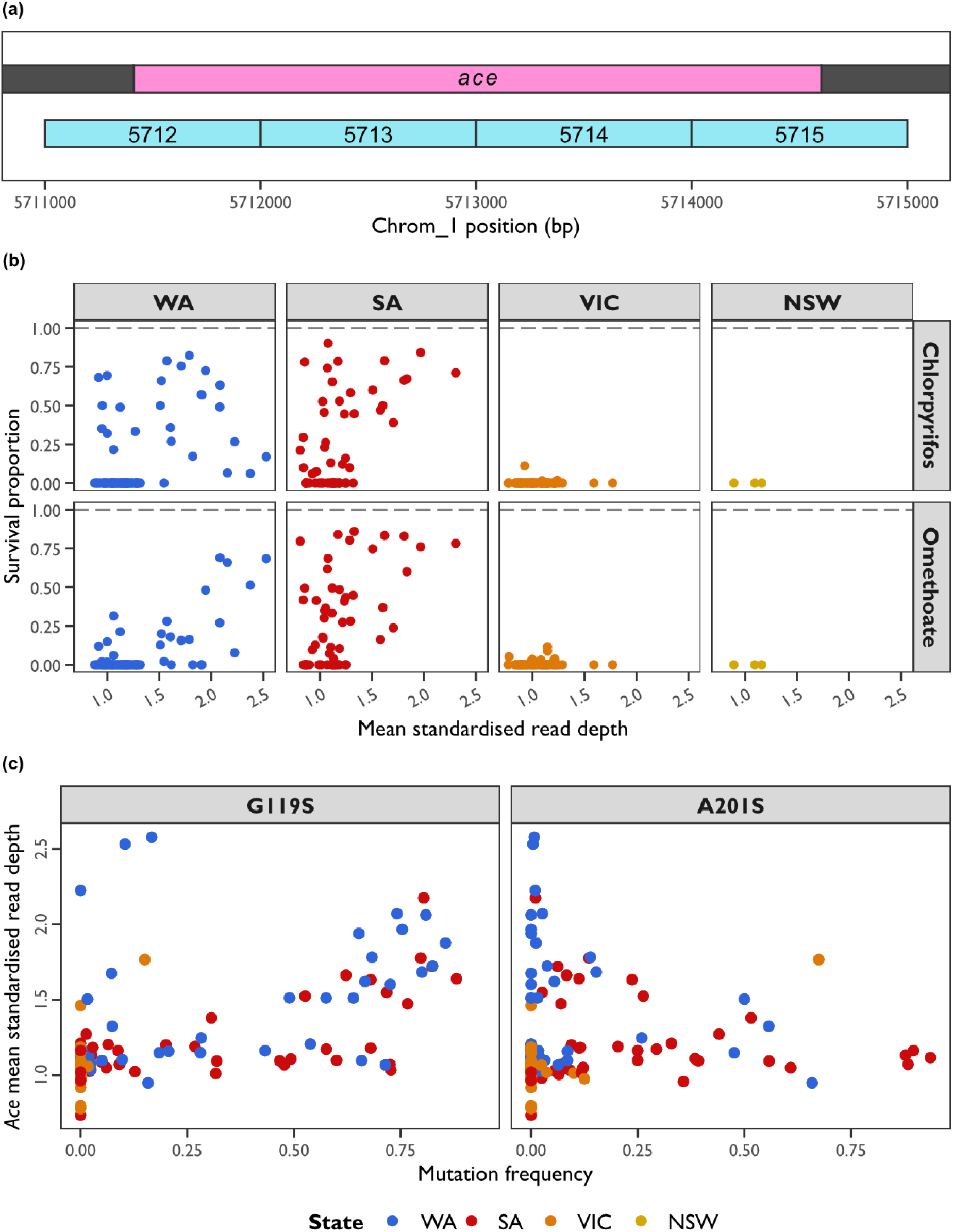
Putative copy number variation in the canonical *ace* gene assessed from standardised read depth across populations of *Halotydeus destructor*. (a) Schematic of the *ace* gene (pink) relative to the 1,000 bp genomic windows analysed (blue). (b) *Ace* copy number relative to pesticide survival. The *x*-axis represents the mean standardised read depth across windows, and the *y*-axis represents the pesticide survival proportion. Points represent individual populations, coloured by state (see legend). Panels separate data for each pesticide (rows) and genomic window (columns). See also individual models for each combination of pesticide and state in Supplementary Table S2. (c) Associations between target-site mutations and copy number. The *x*-axis represent the mutation frequency, and the *y*-axis represents the mean standardised read depth. Points are coloured by state (see legend). Panels separate data for the G119S and A201S mutations.

Closer examination of the relationship between copy number and mutation frequency may partially explain why our results do not fully align with expectations (Figure 5c). There appears to be a potential positive association between the frequency of the G119S mutation and the copy number of *ace* for a subset of populations. This suggests that in some populations, individuals carry multiple copies of *ace* with G119S. This prevents our models from assessing the significance of read depth independent of the effects of candidate mutations.

Finally, we did not find any cross-mapping between reads mapped to the canonical *ace* gene to other carboxylesterases with *ace*-like homology. This was based on our assessment using a sample with high estimated copy number (RLEM202). Our results suggest that whilst having *ace*-like homologies, these carboxylesterases are sufficiently divergent at the sequence level such that they do not interfere with estimates of copy number (see also Results Section ‘Expansion of *ace*-like carboxylesterase genes’).

### KASP marker development

Based on our results, we developed KASP markers for resistance screening. We used the *ace* G119S mutation to screen for chlorpyrifos resistance, which involves mutation of the GGC wild-type codon to an AGC resistant codon. We used the L1024F mutation to screen for bifenthrin resistance, which involves two different mutations: TTG is the wild-type codon, with TTT being the more abundant (Type 1) resistant codon, and TTC being the less abundant (Type 2) resistant codon. Whereas only a single primer mix was needed to assay the *ace* G119S mutation (KBS-2300-00-436054101), we needed two separate primer mixes for the *para* L1024F mutations, one for the Type 1 resistant codon (KBS-2300-00-436067771) and another for the Type 2 resistant codon (KBS-2300-00-436067772). This meant that two separate assays needed to be run for the L1024F mutation per sample.

We tested our KASP primers on 90 populations that represented a range of populations from the states of WA (*n* = 35), SA (*n* = 39), VIC (*n* = 15), and NSW (*n* = 1) with different levels of resistance to chlorpyrifos and bifenthrin (including pesticide-susceptible populations). Assays were run on a LightCycler using a protocol designed to call diploid genotypes: homozygous wild-type, heterozygous, homozygous mutant. As we were working with pooled DNA samples, these calls were not taken to represent true genotypes, but would indicate the strength of signal for susceptible wild-type (FAM fluorescence) vs resistant mutant alleles (HEX fluorescence), which we used as an indicator of their relative frequencies.

The *ace* G119S assay performed well, with strong signals on both the FAM (susceptible wild-type) and HEX (resistant mutant) channels, such that samples were clearly discriminated from negative controls (Figure 6a). The *para* L1024F mutations performed moderately well, with somewhat weaker FAM and HEX signals (Figure 6a). Whilst most samples tested for L1024F could be discriminated from the controls, a few samples did not yield strong signal and were called as negatives.

**Figure 6.**
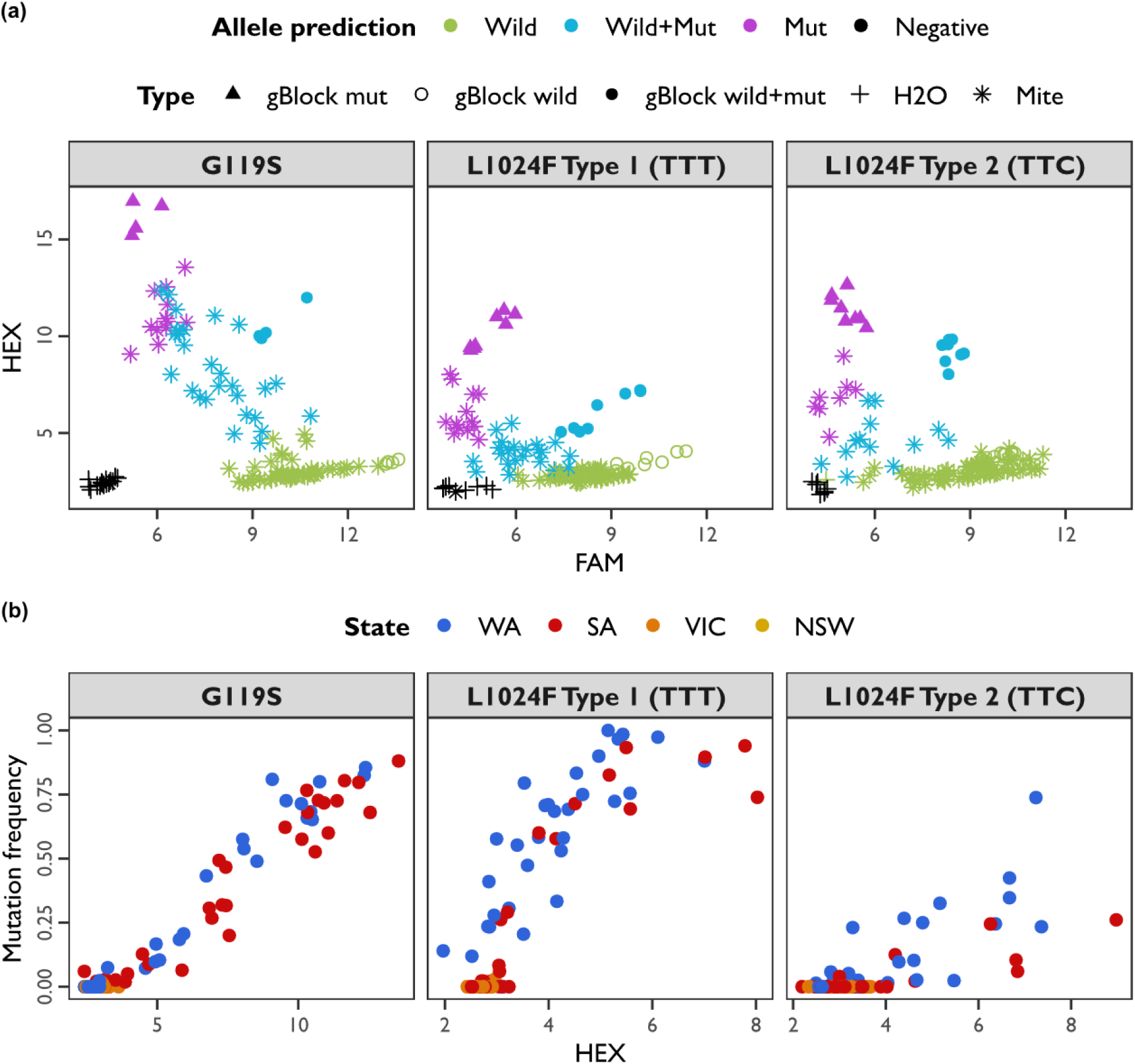
KASP marker application on pooled population DNA samples of *Halotydeus destructor*. (a) Separation of samples along two fluorescence channels, the FAM signal (susceptible wild-type allele) on the *x*-axis, and the HEX signal (resistant mutant allele) on the *y*-axis. Points are coloured based on their genotype call by the LC internal algorithm (see legend); but note that this should not be strictly interpreted as a genotype because these are pooled population samples. Point shapes indicate the type of sample (see legend). Samples include synthesized gBlocks as positive controls (see Main text). (b) Associations between HEX fluorescence on the *x*-axis (note differences in scale) and the resistance mutation frequency on the *y*-axis. Points represent pooled mite population samples, coloured by Australian state (see legend). (a, b) Panels separate mutations (*ace* G119S, *para* L1024F Type 1, and *para* L1024 Type 2).

The HEX signal exhibited strong positive correlations with the pool-seq allele frequencies for the *ace* G119S mutation (*r* = 0.97), and the *para* L1024F Type 1 (*r* = 0.85) and Type 2 (*r* = 0.75) mutations (Figure 6b). The relationships between the HEX signal and allele frequency varied across mutations. Logistic regression slopes were shallowest for the *ace* G119S mutation (*b*_1_ = 0.60), steepest for the *para* L1024F Type 1 mutation (*b*_1_ = 1.74), and comparably shallower for the *para* L1024F Type 2 mutation (*b*_1_ = 0.82). Hence, whilst HEX signals did provide a good reflection of the pool-seq allele frequencies, their relationship varied among mutations.

### Expansion of *ace*-like carboxylesterase genes

We explored the expansion of carboxylesterases with *ace*-like homology by assembling draft genomes from mites sampled in the populations RLEM102 (susceptible) and RLEM120 (omethoate and bifenthrin resistant). The Nanopore long read assemblies produced using *flye2* were 65.4 Mb (RLEM102) and 84.5 Mb (RLEM120) in size. Note that these were not curated as comprehensively as our focal reference genome. BUSCO scores for RLEM102 were Core:64.0% [Single:59.3%, Duplicate:4.7%], Fragmented:3.4%, Missing:32.6%, indicating a generally incomplete assembly. But for RLEM120, BUSCO scores were Core:87.1% [Single:79.2%, Duplicate:7.9%], Fragmented:2.9%, Missing:10.0%, indicating a genomic completeness similar to our focal reference assembly.

Following annotation, and filtering for large proteins ≥400 amino acid residues, there were 18 *ace*-like genes in the assembly of RLEM102, but the canonical *ace* was not recovered. For RLEM120, we observed 23 *ace*-like genes, and the canonical *ace*. For comparison, there were 22 *ace*-like genes that were ≥400 amino acid residues in our focal reference assembly. Our alignment comprised a total of 63 protein sequences with 229 informative sites (amino acids that were polymorphic and excluding gaps).

Our protein alignment indicated potential variation in the composition of *ace*-like carboxylesterases across populations (Figure 7). Not all genes observed in the focal reference genome had homologues in RLEM102 and (or) in RLEM120. Similarly, there were also genes observed in both RLEM102 and RLEM120 not present in the focal reference genome. This pattern is potentially driven by two factors: true variability in the genic content among assemblies, or different assemblies capturing unique non-overlapping pieces of the *H. destructor* genome.

**Figure 7.**
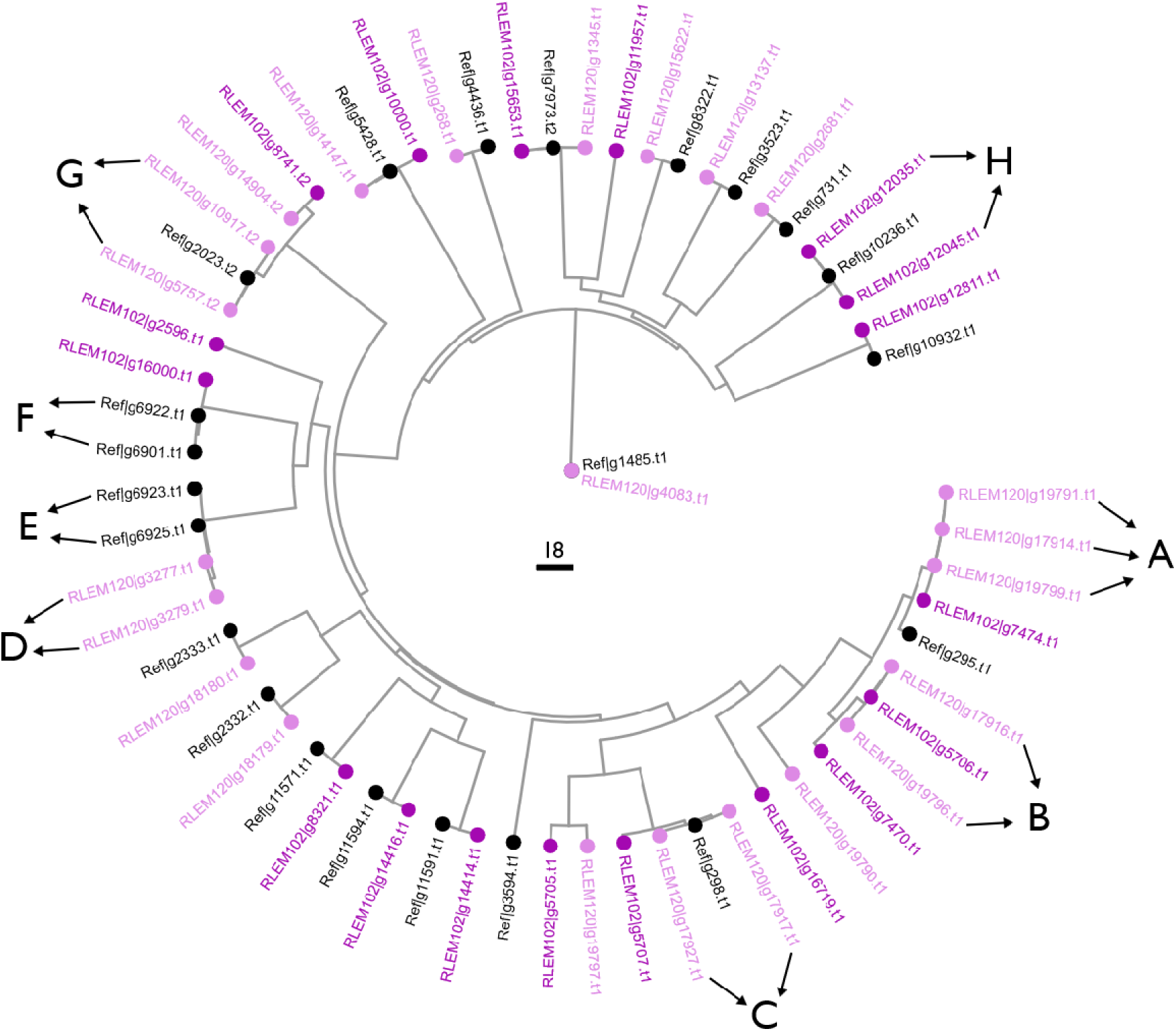
Protein distance tree for annotated *ace*-like genes in the focal *Halotydeus destructor* reference genome and two additional assemblies derived from distinct field populations. The tree was derived from 250 informative amino acid residues. Black points indicate *ace*-like proteins found in the focal reference assembly. Light and dark purple points indicate *ace*-like proteins found in the assemblies from two field populations: RLEM102 and RLEM120. The tree is rooted on the reference canonical *ace* protein, ‘Ref|g1485.t1’. The canonical ace was recovered in the assembly from RLEM120, ‘RLEM120|g4083.t1’, but not from the assembly of RLEM102. Arrows pointing to uppercase letters indicate highly similar genes grouped into clusters. See also Supplementary Figures S6 to S13 for the genomic landscape around genes in these clusters.

We observed 8 groups of genes that exhibited high similarity within their sampled population. At the informative sites, genes within clusters A, D, F, and H exhibited no amino acid differences. The pair of genes in cluster B exhibited 1 amino acid differences, and the pair of genes in cluster C, E, and G exhibited 4 amino acid differences. Clusters split on across separate contigs included clusters A, B, G and H (Supplementary Figures S6, S7, S12, & S13), whereas clusters C, D, E and F were not split across different contigs (Supplementary Figures S8 to S11). Clusters split across different contigs tended to have a similar genomic landscape (gene content and arrangement). This is potentially indicative of orthologous copies within a genome existing on different haplotypes that vary in their genomic landscapes.

## DISCUSSION

### An improved reference genome for *Halotydeus destructor*

The original draft genome assembled for *H. destructor* comprised 132 contigs with a total size of 48.9 Mb (Thia, Korhonen, et al., 2023). Using Hi-C sequencing, we elevated that original draft assembly to produce a new chromosome-level assembly, which comprises 9 chromosomes, totalling 45.9 Mb, with 57 unplaced contigs remaining, totalling 3.0 Mb. This chromosome-level assembly is non-redundant based on self-to-self alignment and was complete based on BUSCO scores (90.5% of Arthropod single-copy orthologs were captured by our assembly). These BUSCO scores were similar to a chromosome-level assembly for *T. urticae* (which contained 90.6% of Arthropod single-copy orthologs). This suggests that our *H. destructor* genome has comparable completeness to others assembled for mites.

Comparative analysis between *H. destructor* and *T. urticae* revealed very limited synteny between these species. The lack of synteny suggests considerable genomic divergence between these mite species. *Halotydeus destructor* belongs to one of the most ancestral families of trombidiform mites, the Penthaleidae, whereas *T. urticae* is in one of the most derived, the Tetranychidae (Klimov et al., 2018; Qin, 1996; Szudarek-Trepto et al., 2022). Chromosome numbers vary markedly across trombidiform mites, ranging from as few as 1 to as many as 14 (Helle et al., 1972; Helle & Bolland, 1967; Shirai et al., 1984). This hints at the potential evolvability in the genomic structure of trombidiform mites, but to date, there have been few comparative analyses in this Order—in part, perhaps, because genome assemblies have been scarce. Our chromosome-level assembly therefore expands the available high-quality genomic resources for comparative analyses of the Trombidiformes.

### Population genetic structure across Australian populations

Our large population genomic dataset has provided a clearer picture of the population structure of *H. destructor* in Australia. Prior studies have been constrained in their scope in terms of geographic scale, the number of populations sampled, the continuity of sampling across space, and (or) genetic markers used (Hill et al., 2016; Thia, Korhonen, et al., 2023; Weeks et al., 1995; Yang et al., 2020). In the present study, sampling occurred across a large part of the Australian distribution of *H. destructor*, capturing a continuous range of distances both within and between western and eastern regions. Whereas previous studies have supported clear west–east breaks in genetic structure (Thia, Korhonen, et al., 2023; Yang et al., 2020), our new analyses reveal a more nuanced pattern. Genetic variation exhibits a continuum between being ‘west-like’, populations from WA, and ‘east-like’ populations from VIC and NSW, with SA populations somewhere along this continuum. This makes sense given the historical expansion of *H. destructor*, which initially invaded Australia in the west, and then spread into eastern Australia during the 1920s (Ridsdill-Smith, 1997). This serial invasion pathway is also captured in patterns of genetic diversity, which exhibited an overall west-to-east decline, and corroborates findings made using a much smaller sample size (Thia, Korhonen, et al., 2023).

Although broad west-east genetic structuring was evident in the present study, there was also a more complex spatial organisation of genetic variation among populations of *H. destructor*. Nested within the regional structuring of genetic variation was an isolation-by-distance pattern. Such hierarchical patterns of structure can emerge when different population genetic processes operate at different spatiotemporal scales (Kamel et al., 2014; Meirmans, 2012). The patterns of isolation-by-distance were logarithmic, in that *F*_ST_ rose quickly over scales of tens of kilometers but plateaued or became more variable on the scale of hundreds to thousands of kilometers. These results are in line with previous studies investigating genetic structure on the scale of hundreds of kilometres (Weeks et al., 1995), and on the finer scale of hundreds of meters (Yang et al., 2020). Studies at more localised scales have also found evidence of very fine microgeographic isolation-by-distance across tens of meters (Yang et al., 2020), but we did not sample at such scales here.

Finally, patterns of isolation-by-distance themselves exhibited spatiotemporal heterogeneity, as evidenced by differences in the slope within west vs east regions and the variance in this relationship. In other organisms, spatiotemporal heterogeneity in environmental and demographic factors can cause variable patterns in genetic structure (Gould & Dunlap, 2017; Levy & Neal, 1999; Thia, McGuigan, et al., 2021). Teasing out such factors was not the goal of this work, but remains of interest for future investigation.

### Target-site mutations and their associations with pesticide resistance

Our population genomic data allowed us to characterise the allelic variation in the target genes of three important chemical pesticides used to control *H. destructor*: the *ace* (acetylcholinesterase), *para* (voltage-gated sodium channel) and the *nAChR* (nicotinic acetylcholine receptor) genes. Although there was an appreciable amount of polymorphism within these genes, many of the non-synonymous mutations were rare (below the minor allele threshold of 0.05). Our association tests focused on 3 common non-synonymous mutations: G119S and A201S (*T. californica* numbering) in the *ace* gene, and L1024F (*M. domestica* numbering) in the *para* gene. These three mutations have previously been described in *H. destructor* (Edwards et al., 2018; Thia, Korhonen, et al., 2023; Yang et al., 2020). The two *ace* mutations underpin resistance in other arthropod pests (Carvalho et al., 2013; Khajehali et al., 2010), and a similar *para* mutation, L1024V, contributes to resistance in *T. urticae* (Riga et al., 2017).

Our genotype-phenotype associations for the *ace* target-site mutations revealed some interesting patterns. Whilst G119S showed a clear positive correlation with chlorpyrifos resistance, the association with omethoate resistance was not clear. There was also no clear association between A201S and resistance to either organophosphate compound. These data suggest that the genetic architecture of resistance for chlorpyrifos and omethoate may be different (at least partially) such that there is a low genetic covariance (low cross-resistance) between them. The impact of A201S on resistance may depend on genetic background, as shown in *T. urticae* across different pesticides (Riga et al., 2017). While A201S likely contributes to organophosphate resistance in some cases, its effect seems weaker and more variable than that of G119S. We also note that A201S and G119S show an unusual frequency pattern. Many populations are fixed for one of the resistance mutations but lack the other one, whereas populations that are polymorphic for both mutations exhibit a negative correlation between them (Figure S4a). This mirrors our earlier findings based on fewer populations (Thia, Korhonen, et al., 2023) and suggests that selection for G119S may reduce the frequency of A201S, or vice versa (Thia, Korhonen, et al., 2023).

Our data are consistent with the hypothesis that chlorpyrifos resistance may predictably evolve across populations as a monogenic trait underpinned largely by the G119S target-site mutation. In contrast, other genetic mechanisms may play a more important role in omethoate resistance. The biochemical properties of these compounds differ: chlorpyrifos contains diethyl groups attached to its phosphate moiety, whereas omethoate contains dimethyl groups. Typically, organophosphates with diethyl groups are more toxic than those with dimethyl groups and have greater inhibitory capacity of other serine esterases (Chambers & Chambers, 1989; Meek et al., 2021). These may select for different resistance mechanisms between chlorpyrifos and omethoate, resulting in low cross-resistance. Copy number variation in *ace* might also play an important role (see below).

The *para* L1024F mutation exhibited a strong positive correlation with bifenthrin resistance. However, the slope of the relationship was less than 1 (Figure 3). These findings contrast with observations from populations collected in 2014 and 2015, where the correlation between the L1024F mutation and bifenthrin resistance approximated a slope of 1 (Edwards et al., 2018). One possible explanation for this apparent shift may be due to alternative mechanisms for pyrethroid resistance. It has been nearly two decades since pyrethroid resistance first appeared in Australian populations of *H. destructor* (Umina, 2007). Since then, increasing evidence suggests that target-site resistance to pyrethroids comes at a cost, and this may be partially mediated by trade-offs with climatic adaptation (Maino et al., 2018; Umina et al., 2024). Indeed, studies in other arthropods have shown that *para* target-site mutations can come with a range of pleiotropic effects with negative fitness consequences (Foster et al., 2002; Freeman et al., 2021).

If fitness costs of the L1024F mutation are strong, mutations that introduce genetic redundancy in pyrethroid resistance may be favoured (Láruson et al., 2020). Prior work did suggest that other metabolic mechanisms may partially contribute to resistance (Yang et al., 2020). However, our present work also shows evidence of other target-site mutation segregating in the *para* gene (see below). Whilst we expect the *para* L1024F mutation to still be important in resistance at broad spatial scales (across populations), other target-site mutations might make important contributions at more localised spatial scales (within populations). It is also possible that there are increasing roles of metabolic mechanisms evolving in response to pyrethroid pesticides (Yang et al., 2020).

A final point is that our study revealed a number of known target-site mutations that were rare and below the minor allele threshold of 0.05. For the *ace* gene, this included the F331Y mutation, which is known from previous work (Thia, Korhonen, et al., 2023) and has been associated with organophosphate resistance in other mites (Carvalho et al., 2012; Khajehali et al., 2010). For the *para* gene, 5 other target-site mutations were observed. M918V and M918T are the classic mutations associated with *super-kdr* phenotypes in other hemipterans (Eleftherianos et al., 2008; Morin et al., 2002). L925I has been linked to resistance in silverleaf whitefly, *Bemisia tabaci* (Morin et al., 2002), and has been observed previously in *H. destructor* (Thia, Korhonen, et al., 2023). F1538I has been linked to resistance in two-spotted spider mite, *T. urticae* (Tsagkarakou et al., 2009). The detection of *para* M918V, M918T, and F1538I is the first record of these mutations in *H. destructor*. An additional fifth rare mutation that we observed in this work was the *para* F1020Y, which we have documented previously (Thia, Korhonen, et al., 2023), but no other study has linked this specific amino acid substitution to resistance. However, a similar mutation, F1020S, has been shown to contribute to resistance in diamondback moth, *Plutella xylostella* (Endersby et al., 2011).

### Copy number variation in *ace* has mixed associations with organophosphate resistance

Our analyses of mean copy number did not provide consistent support that increased copies of *ace* are associated with organophosphate resistance in *H. destructor*. Amplification of *ace* has been observed in other arthropods, conferring resistance by effectively diluting the pesticide relative to the number of enzyme molecules that must be inhibited to achieve mortality (R. Carvalho et al., 2012; Kwon et al., 2010). However, a nuance to this result is that the effects of target-site mutations and copy number in *ace* cannot fully be disentangled with our current pool-seq data. We found that a subset of *H. destructor* populations appear to show a positive association between the frequency of the G119S mutation and estimated *ace* copy number. Amplification of *ace* copies carrying target-site mutations has been observed in spider mites (R. Carvalho et al., 2012; Kwon et al., 2010) and in mosquitoes (Alout et al., 2009).

The joint polymorphism of copy number and target-site mutations within and among populations of *H. destructor* increases the complexity of organophosphate resistance at the broad-scale. Our data hints at the possibility of many segregating haplotypes in the genomic region containing *ace*, involving different numbers of *ace* gene copies that themselves carry different compositions of alleles. Individual-level sequencing will be required to tease out these genetic contributions. However, a key result from this work was that the pooled frequency of the G119S mutation is strongly correlated with chlorpyrifos resistance. Hence, irrespective of *ace* copy number, the relative frequency of G119S predictably indicates the prevalence of chlorpyrifos resistance within *H. destructor* populations.

### New KASP markers for genetic monitoring of pesticide resistance

We developed *ace* G119S and *para* L1024F as KASP markers for chlorpyrifos and bifenthrin resistance, respectively, given their strong associations with resistance to these pesticides. We applied the KASP markers to pooled DNA extractions because this provides the most efficient use-case in which a single sample can be screened for resistance and quantified for the frequencies of causal alleles. The feasibility of this approach has been demonstrated previously, for example, when screening for resistance in other arthropod pests (Shi et al., 2023), and when screening for rare alleles in seed mixtures (Brusa et al., 2023).

Our *ace* G119S KASP assay comprises a single primer set to distinguish between the G119 and S119 alleles, a GGC wild-type codon and an AGC resistant codon, respectively. Amplification of these G119S primers was strong and there was a tight positive relationship between the HEX signal (mutant S119 allele) and the allele frequency estimated using whole-genome pool-seq. Our *para* L1024F KASP assay comprises a set of two primer sets. The first set is used to distinguish between the L1024 and Type 1 F1024 alleles, a TTG wild-type codon and a resistant TTT codon, respectively. The second set is used to distinguish between the L1024 and Type 2 F1024 alleles, the resistant allele this time being the rarer TTC codon. Amplification of these L1024F primers was moderate, and we suspect that intraspecific variation, especially at the same nucleotide position, may reduce the overall efficiency of the assay. Nonetheless, the HEX signal (mutant Type I and 2 F1024 alleles) still exhibited a positive relationship with expected whole-genome pool-seq allele frequencies.

Our new KASP markers expand available molecular tools for genetic monitoring of resistance in *H. destructor* (Edwards et al. 2018; Cheng et al., 2019; Umina et al. 2024). For applications to pooled DNA samples, the in-built genotype calling protocol in LightCycler instruments has the utility of binning samples into low (homozygous wild-type), medium (heterozygous), and high (homozygous mutant) risk in terms of potential underlying mutant allele frequencies. For quantitative estimation, we suggest that studies perform preliminary runs using samples with known allele frequencies to determine the relationship with HEX signal under local laboratory conditions and platforms. Once this relationship is established, these KASP markers can be effectively used to screen pooled DNA samples of unknown allele frequencies.

### Carboxylesterases with *ace*-like homology are potentially diverse within and among populations

We used long-read data to explore the variation in carboxylesterase genes that exhibited *ace*-like annotations. In prior work, we have shown that the *H. destructor* genome has a large number of carboxylesterases that exhibit homology to the canonical *ace* gene (Thia, Korhonen, et al., 2023). These *ace*-like carboxylesterases appear to exhibit copy number variation within and among populations, and there is some evidence that collectively these genes might play some role in organophosphate resistance (Thia, Umina, et al., 2023). Indeed, it is well known that carboxylesterases are often co-opted into pesticide metabolism through increased expression or amino acid changes in other arthropod pests (Field, 2000; Newcomb et al., 1997; Thia et al., 2024).

Here, the variation we observed in the *ace*-like carboxylesterases took on different forms. First, a single orthologous gene might exist in different haplotypes that capture different genomic landscapes around them. Second, closely related paralogous genes might coexist in the same genome, representing potential gene duplication events. Thirdly, different genomes might vary in their composition of these genes. Our understanding how these different *ace*-like carboxylesterases contribute to pesticide adaptation in *H. destructor* is still in its infancy. Pangenomic approaches might provide a useful path forward (Jiao & Schneeberger, 2020; Kang et al., 2023). Such approaches would provide greater scope for characterising how these genes and associated haplotypes segregate within and among populations, and a means to link this variation to pesticide resistance.

### Conclusions and outstanding questions

High-quality reference genomes and spatiotemporally replicated population genomic datasets are key resources for developing genetically informed resistance management strategies and monitoring tools (Fritz, 2022; Thia, Hoffmann, et al., 2021). The population genomic analyses in this study advance previous work by illuminating the complex spatial patterns of genetic structure among *H. destructor* populations. Whilst there is evidence of regional structure, and some evidence of isolation-by-distance, other environmental factors may shape the distribution of genetic variation across the broader landscape—particularly within western and eastern regions.

Our data supports previous suggestions that target-site resistances likely evolve repeatedly at more localized scales, instead of being spread by dispersal over large geographic distances (Thia, Korhonen, et al., 2023; Yang et al., 2020). Moreover, our broad-scale population genomic dataset (spanning thousands of kilometers and multiple years) provides the most detailed survey of the frequency and distribution of target-site mutations in *H. destructor* to date. This was facilitated by our pool-seq approach, allowing us to sample many more populations and screen many more genomic positions that what would have been possible using individual-sequencing and locus-specific genotyping (such as Sanger sequencing). Linking this information to phenotypic data was crucial in establishing candidates for KASP markers, which provide a rapid and affordable diagnostic for resistance.

Our KASP markers for chlorpyrifos and bifenthrin resistance are useful because the underlying target-site alleles have strong associations with resistance. But findings also suggest an increasing diversity of target-site mutations, and putative other mechanisms, segregating within and among field populations. The expanding complexity of known resistance mechanisms in *H. destructor* iterates the need for ongoing spatiotemporal genetic data. Such long-term, and spatially broad, datasets are pivotal in tracking shifts in resistance mechanisms (Thia, Hoffmann, et al., 2021). This is key to evaluating the ongoing efficacy of diagnostic molecular markers.

An outstanding issue is how frequently long-distance dispersal occurs and whether this has a significant impact on local population genetic processes. Whilst this issue has been explored previously, the recent invasion history of Australian *H. destructor*, and their large population sizes, have been difficult to disentangle from the effects of gene flow in demographic models (Thia, Korhonen, et al., 2023). Additionally, determining the function of *ace*-like carboxylesterases could help us better understand the role they may play in pesticide adaptation (Thia, Umina, et al., 2023). This could possibly explain some of the unaccounted variation in omethoate resistance that appears poorly associated with *ace* target-site mutations. Both questions remain as areas of interest for future investigations.

## ACKNOWLEDGEMENTS

Genomic and molecular work was supported through an investment from the Grains Research and Development Corporation (UOM1906-002RTX; UOM2404-006RT) and Hort Innovation (ST23002) as part of the ‘Australian Grain and Horticulture Pest Innovation Program’ to PAU and AAH, with additional investment from the University of Melbourne and Cesar Australia. Mite collections and phenotypic testing was made possible through funding from the Grains Research and Development Corporation (UM00057 and CES2010-001RTX), Meat and Livestock Australia (P.PSH.1283), AgriFutures Australia (PRJ-013101) to PAU, and the Australian Research Council (DE230100257) to TLS. This research was supported by The University of Melbourne’s Research Computing Services and the Petascale Campus Initiative. We would also like to thank Adriana Arturi, Paul Tyson, James Maino, Luis Mata and Anthony van Rooyen for technical input, assisting with field collections and molecular screening of pyrethroid resistance.

## AUTHOR CONTRIBUTIONS

JAT and AAH conceived and designed the genomic components of this study. JAT led all bioinformatics, population genomics, and the design of the KASP experiments and analysis; he also contributed to molecular work. KR led molecular work associated with KASP assays. CB contributed to molecular work associated with the population genomic dataset. SBS led molecular work associated with generating Hi-C libraries (Ref assembly) and Nanopore long-reads (RLEM102 and RLEM120 assemblies). NDY, RBG, and BCHC contributed support to the reference genome assembly. SW contributed to the design of the KASP experiments. JC contributed to the long-read analyses. AA, PAU, SM, and AL led the collection of mite samples and undertook the phenotypic testing. PAU and AAH secured funding for collections and population genomic works. TLS contributed funding toward generation of Nanopore long-read data (RLEM102 and RLEM120 assemblies).

## DATA AVAILABILITY

Our chromosome-level reference genome has been submitted to NCBI under accession GCA_022750525.2. All pool-seq Illumina short-read libraries have been deposited under the NCBI BioProject PRJNA756307. The NCBI SRA and BioSample numbers for these pool-seq samples are recorded in Appendix 2. Data and scripts for running analyses have been deposited into a University of Melbourne repository hosted by FigShare (Thia, 2025).

## SUPPLEMENTARY INFORMATION

**Figure S1.**
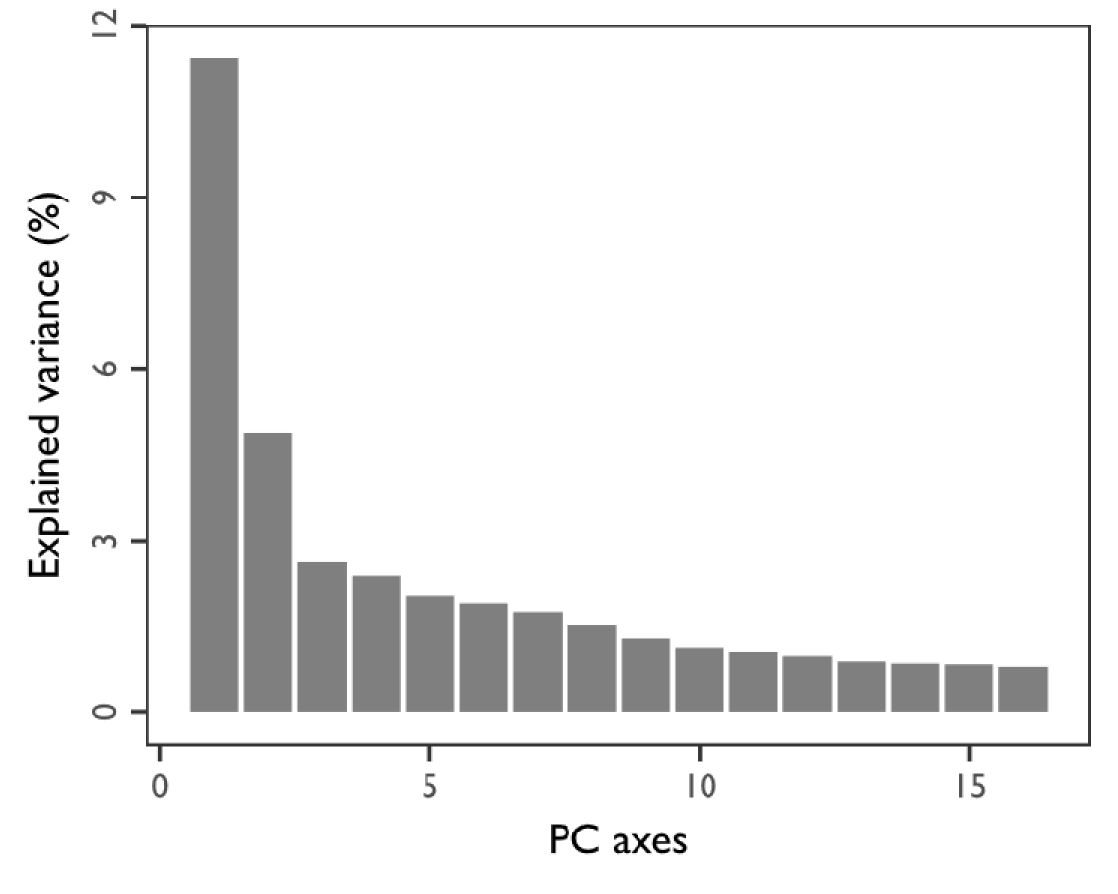
Scree plot for a PCA of nuclear SNP allele frequencies in *Halotydeus destructor*. The *x*-axis represents the first 15 PC axes, and the *y*-axis represents the percentage of explained variance. Note the elbowing at PC3, suggesting that PC1 to PC2 explain most of the population genetic structure.

**Figure S2.**
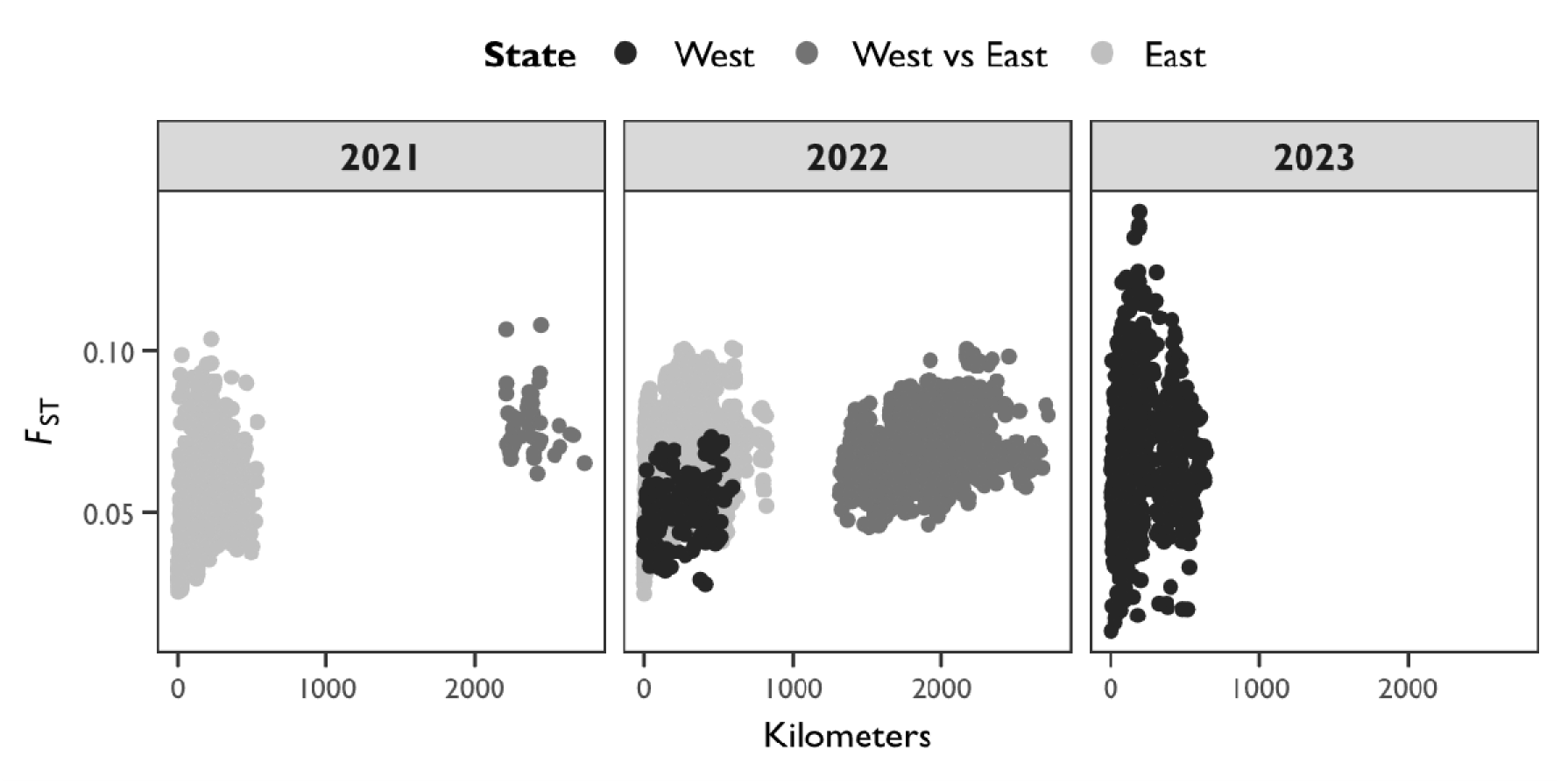
Isolation-by-distance between *Halotydeus destructor* population pairs within Australia. Geographic distance is on the *x*-axis, measured in kilometers. Genetic differentiation is on the *y*-axis, measured as *F*_ST_. Points represent population pairs, coloured by region (west or east) (see legend). Panels separate population pairs from different years.

**Figure S3.**
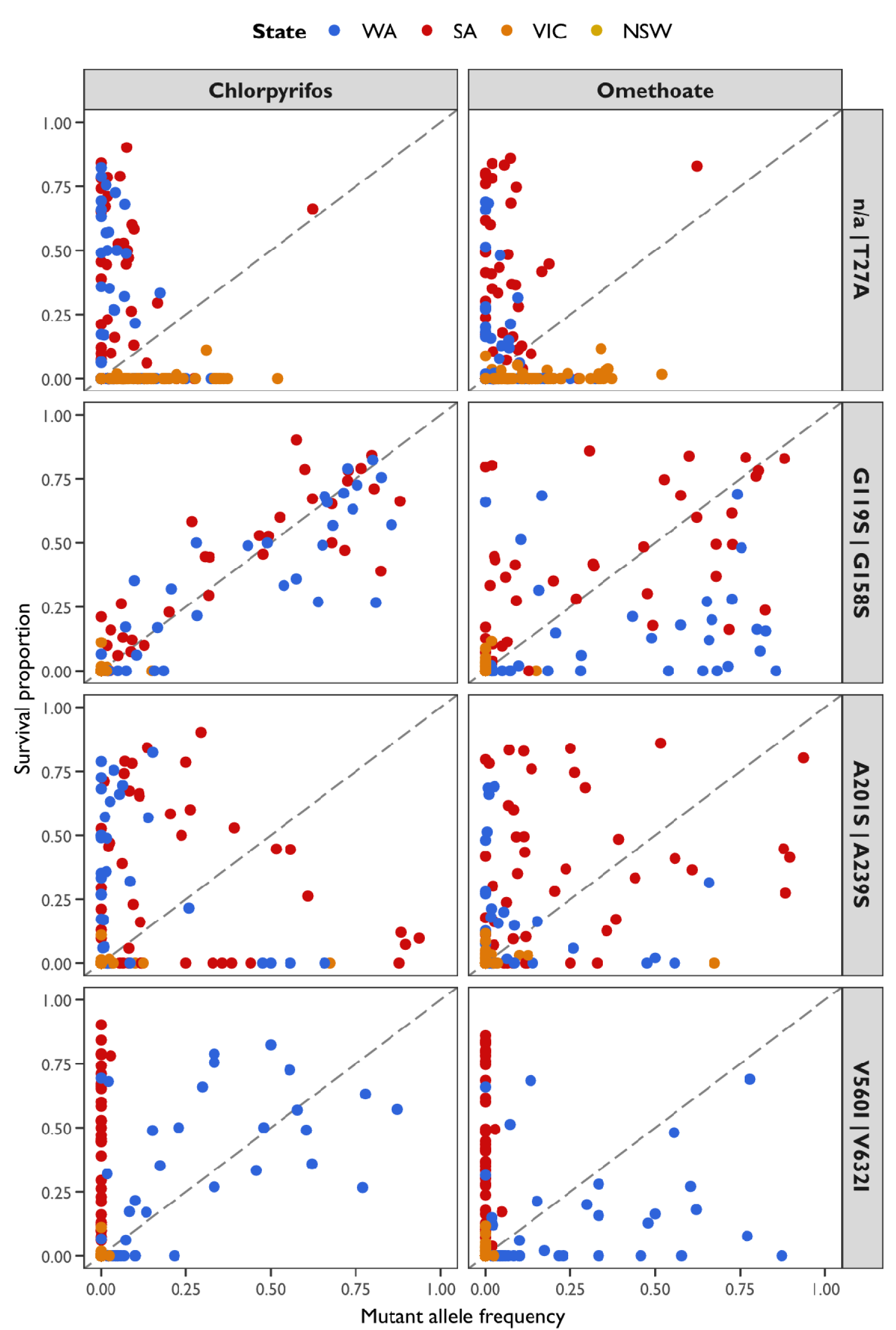
Associations between non-synonymous mutations in the *ace* gene and organophosphate resistance in *Halotydeus destructor*. The *x*-axis represents the allele frequency of each mutation, and the *y*-axis represents the proportional survival of mites following exposure to a discriminating concentration of chlorpyrifos and omethoate. Points represent mite populations, coloured by Australian state (see legend). The dashed lines indicate an expected 1:1 relationship. Panels contain data for combinations of mutation (*Torpedo californica* numbering first, *H. destructor* numbering second, separated by a bar; ‘n/a’ indicates no equivalent position in *T. californica*) and pesticide. See also Supplementary Figure S5 for associations between known target-site mutations G119S and A201S with the mutation V560I: V560I appears to have an association with chlorpyrifos, but is not a known target-site mutation.

**Figure S4.**
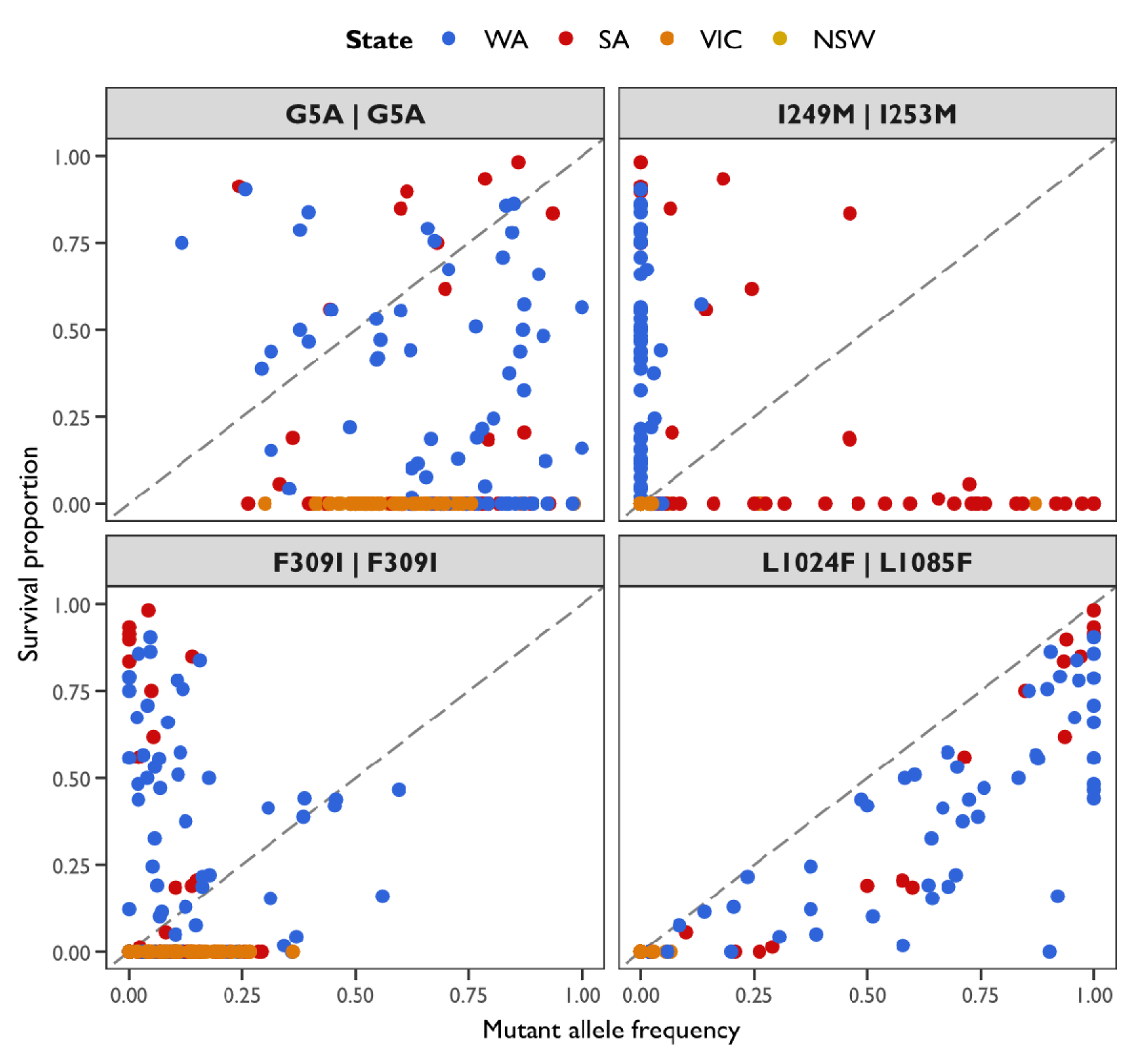
Associations between non-synonymous mutations in the *para* gene and pyrethroid resistance in *Halotydeus destructor*. The *x*-axis represents the allele frequency of each mutation, and the *y*-axis represents the proportional survival of mites following exposure to a discriminating dose of bifenthrin. Points represent mite populations, coloured by Australian state (see legend). The dashed lines indicate an expected 1:1 relationship. Panels contain data for each mutation (*Musca domestica* numbering first, *H. destructor* numbering second, separated by a bar).

**Figure S5.**
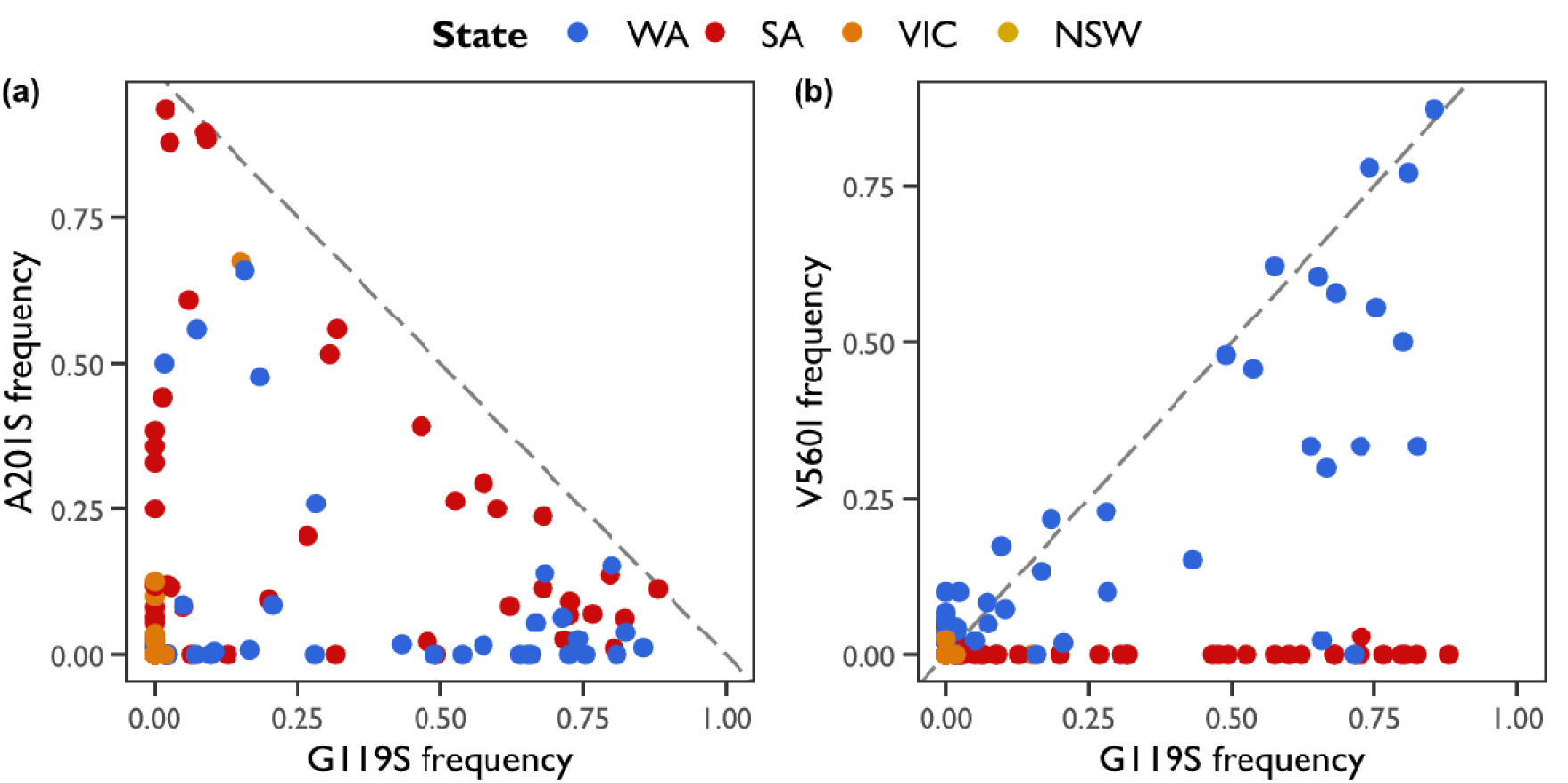
Associations between the *ace* non-synonymous G119S mutation and the A201S and V560I mutations in *Halotydeus destructor*. The *x*- and *y*-axes represent the allele frequency for each mutation (*Torpedo californica* numbering). Points represent mite populations, coloured by Australian state (see legend). (a) G119S vs A201S. The dashed line indicates an expected 1:− 1 relationship. These mutations are uncorrelated in some populations, and negatively correlated in other populations. Negative correlations suggest that in some populations, these mutations are on different haplotypes. (b) G119S vs V560I. The dashed line indicates an expected 1:1 relationship. These mutations are uncorrelated in some populations, and positively correlated in other populations. Positive correlations suggest that in some populations, these mutations are on the same haplotype. Hence, associations between V560I and resistance (as in Figure S3) may be due to its linkage to G119S.

**Figure S6.**
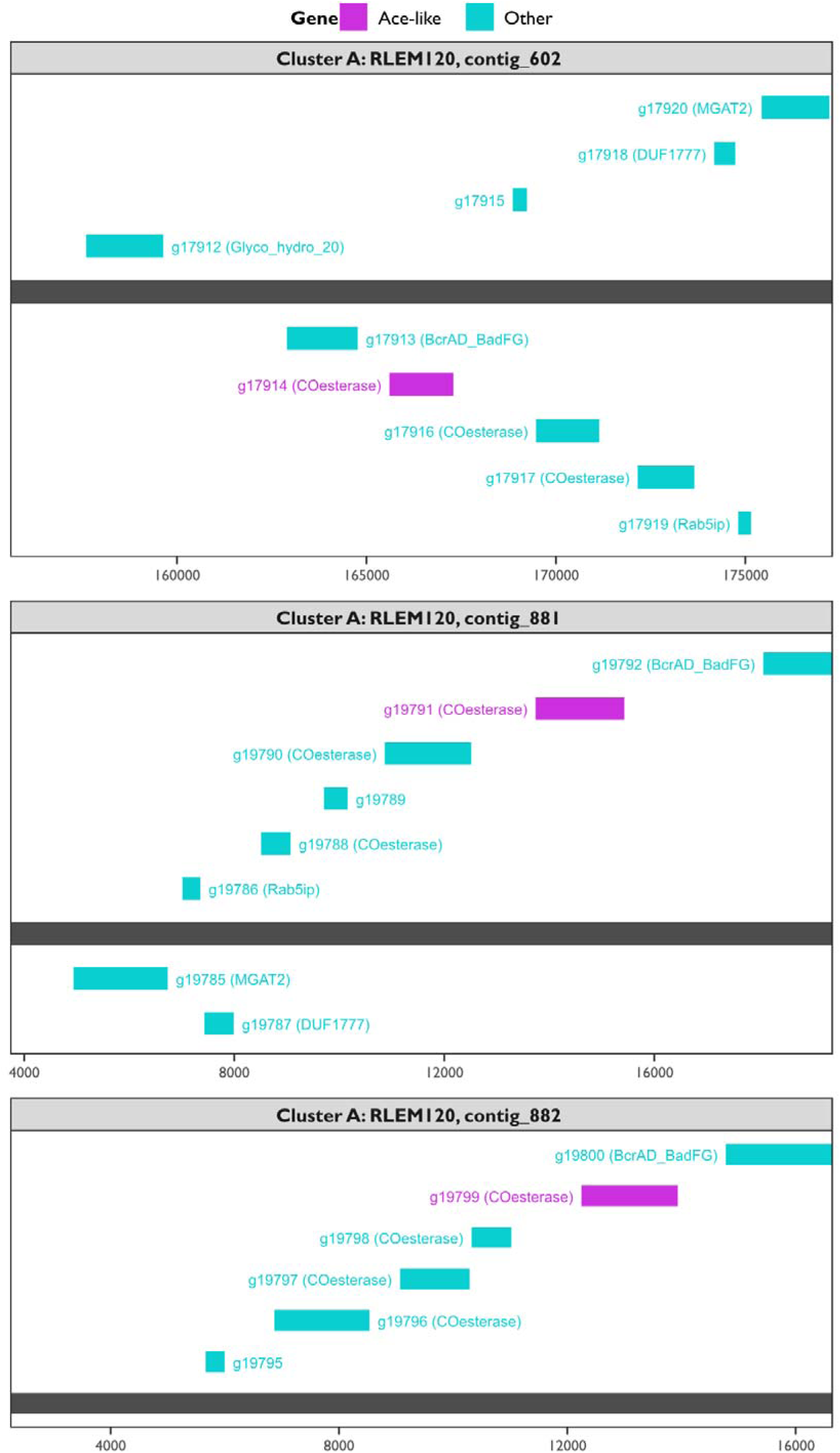
Genomic context around *ace*-like genes in cluster A (as per Figure 7). Data support 3 putative orthologous genes artificially split across different genomic sequences. The *x*-axis represents the base coordinate on the assembled chromosome (or contig). The grey horizontal bar represents the genomic sequence. The magenta bars represent the *ace*-like genes. The turquoise bars represent other genes within 10,000 bp of the *ace*-like genes. Genes *above* the grey genomic sequence are those on the positive strand, whereas those *below* the grey genomic sequence are those on the negative strand. Panels separate different chromosomes (or contigs).

**Figure S7.**
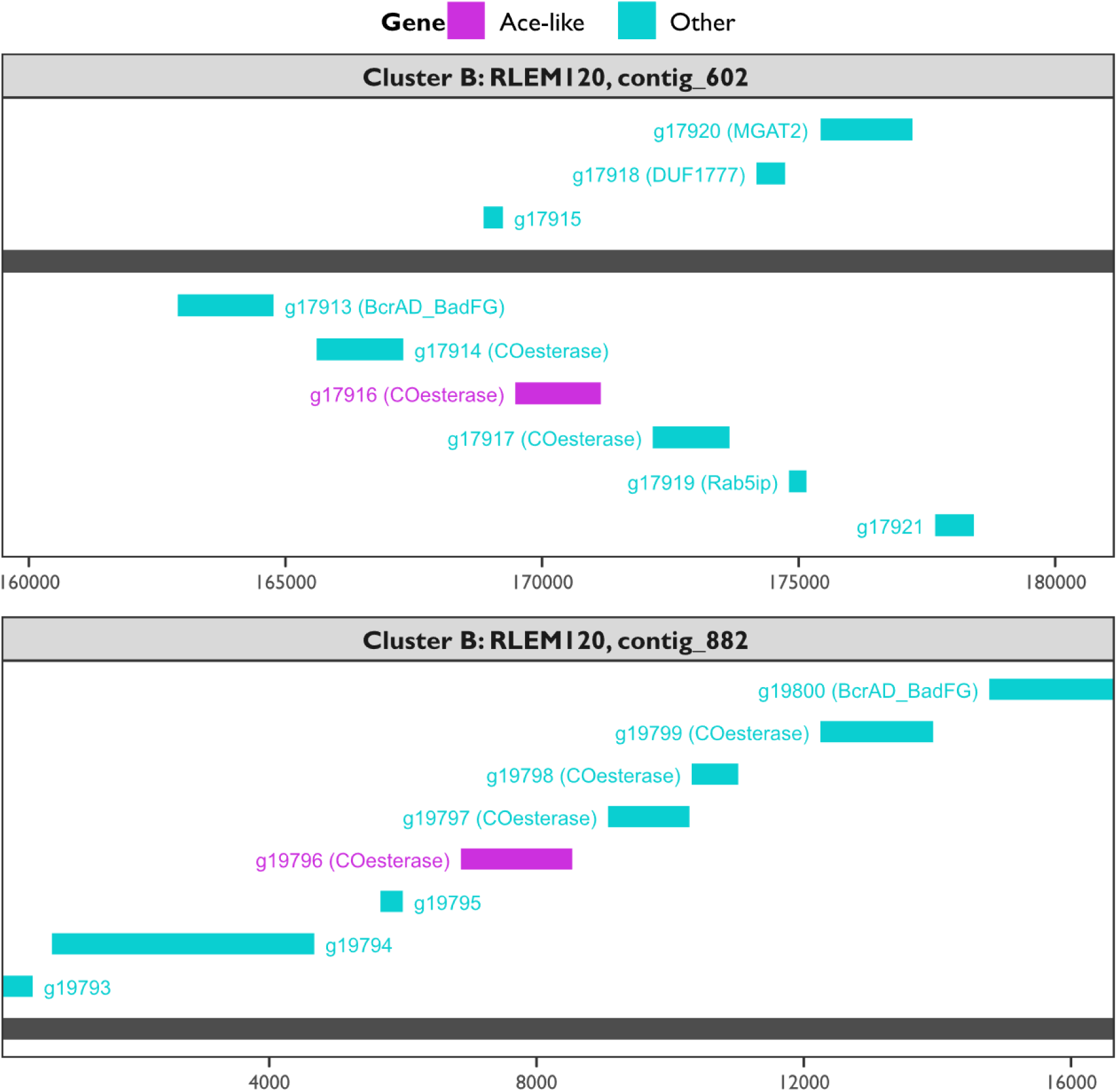
Genomic context around *ace*-like genes in cluster B (as per Figure 7). Data support 2 putative orthologous genes artificially split across different genomic sequences. The *x*-axis represents the base coordinate on the assembled chromosome (or contig). The grey horizontal bar represents the genomic sequence. The magenta bars represent the *ace*-like genes. The turquoise bars represent other genes within 10,000 bp of the *ace*-like genes. Genes *above* the grey genomic sequence are those on the positive strand, whereas those *below* the grey genomic sequence are those on the negative strand. Panels separate different chromosomes (or contigs).

**Figure S8.**
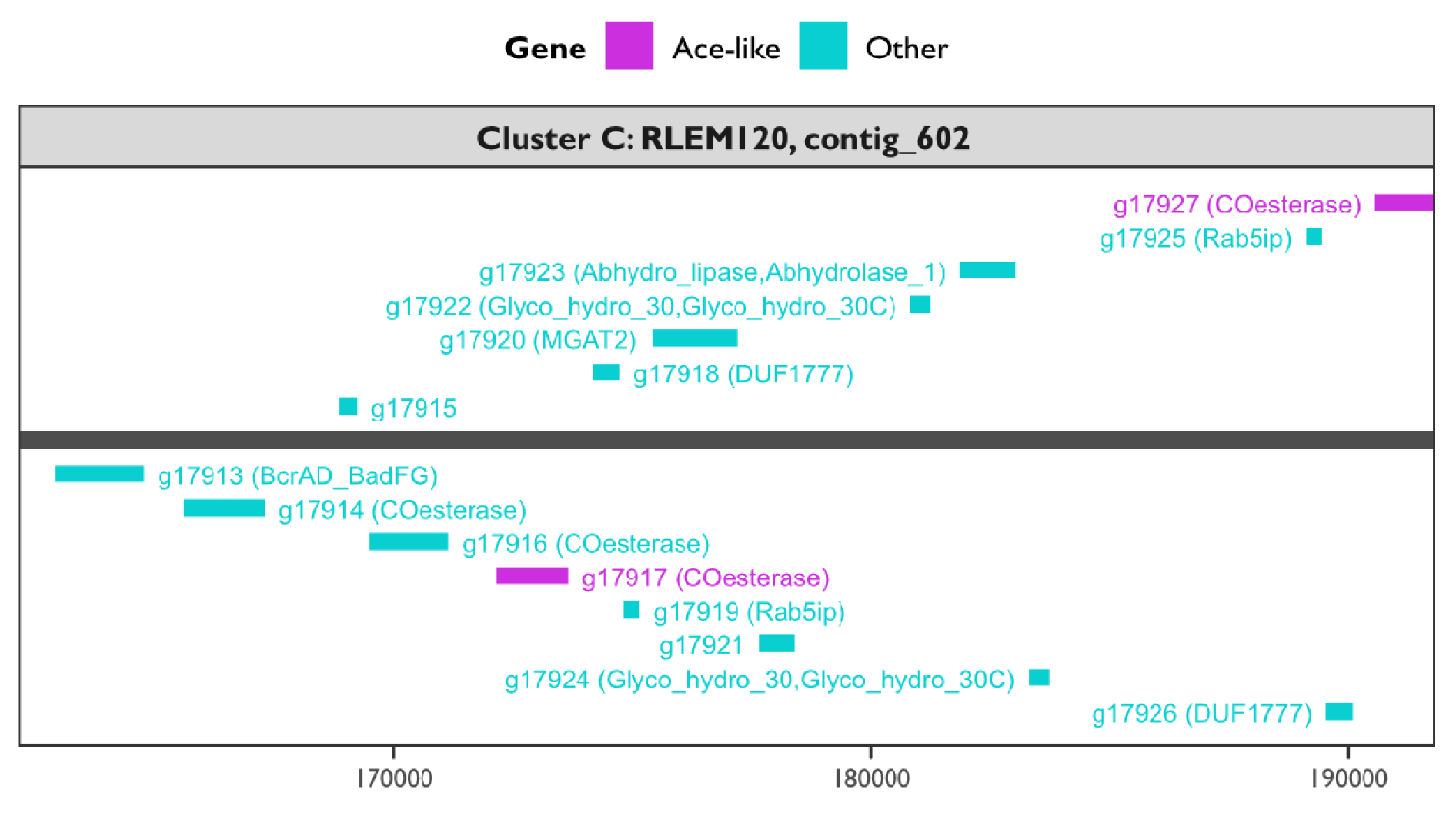
Genomic context around *ace*-like genes in cluster C (as per Figure 7). Data support 2 putative paralogous genes on the same genomic sequence. The *x*-axis represents the base coordinate on the assembled chromosome (or contig). The grey horizontal bar represents the genomic sequence. The magenta bars represent the *ace*-like genes. The turquoise bars represent other genes within 10,000 bp of the *ace*-like genes. Genes *above* the grey genomic sequence are those on the positive strand, whereas those *below* the grey genomic sequence are those on the negative strand. Panels separate different chromosomes (or contigs).

**Figure S9.**
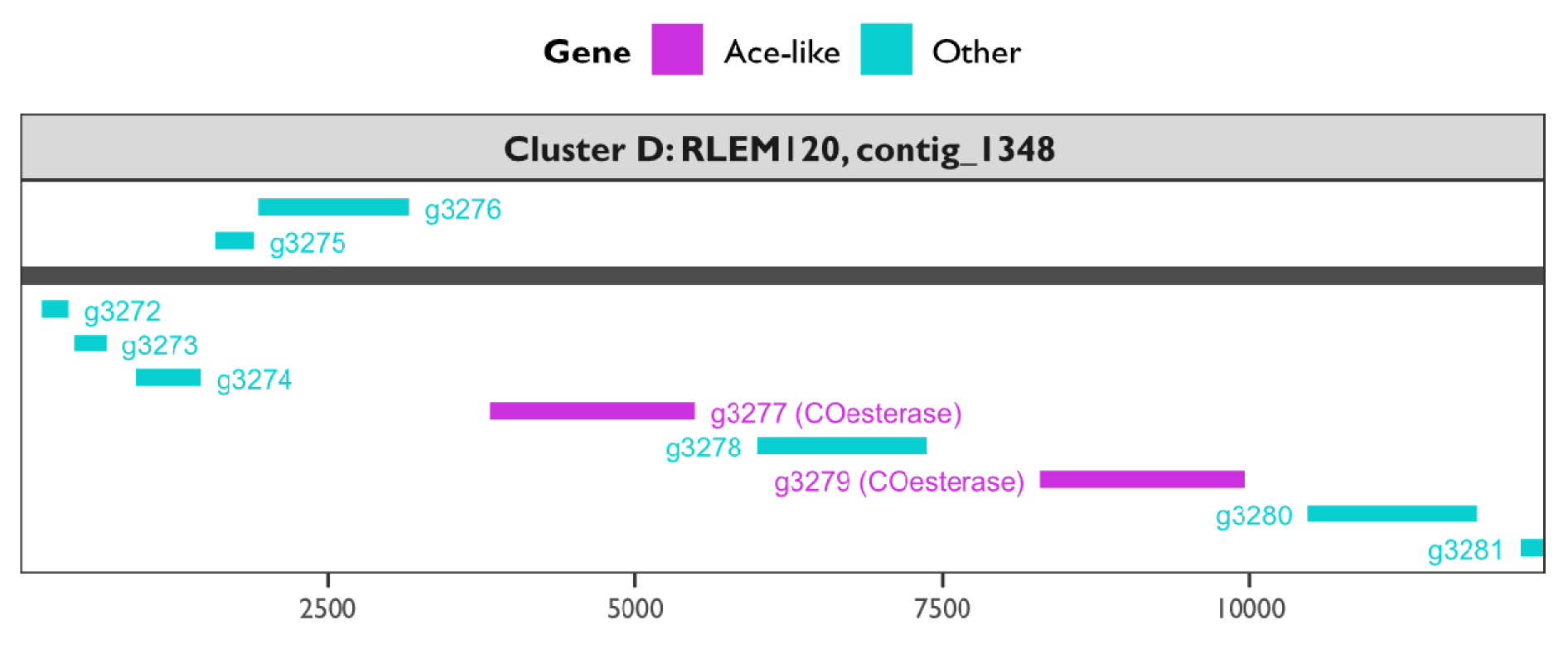
Genomic context around *ace*-like genes in cluster D (as per Figure 7). Data support 2 putative paralogous genes on the same genomic sequences. The *x*-axis represents the base coordinate on the assembled chromosome (or contig). The grey horizontal bar represents the genomic sequence. The magenta bars represent the *ace*-like genes. The turquoise bars represent other genes within 10,000 bp of the *ace*-like genes. Genes *above* the grey genomic sequence are those on the positive strand, whereas those *below* the grey genomic sequence are those on the negative strand. Panels separate different chromosomes (or contigs).

**Figure S10.**
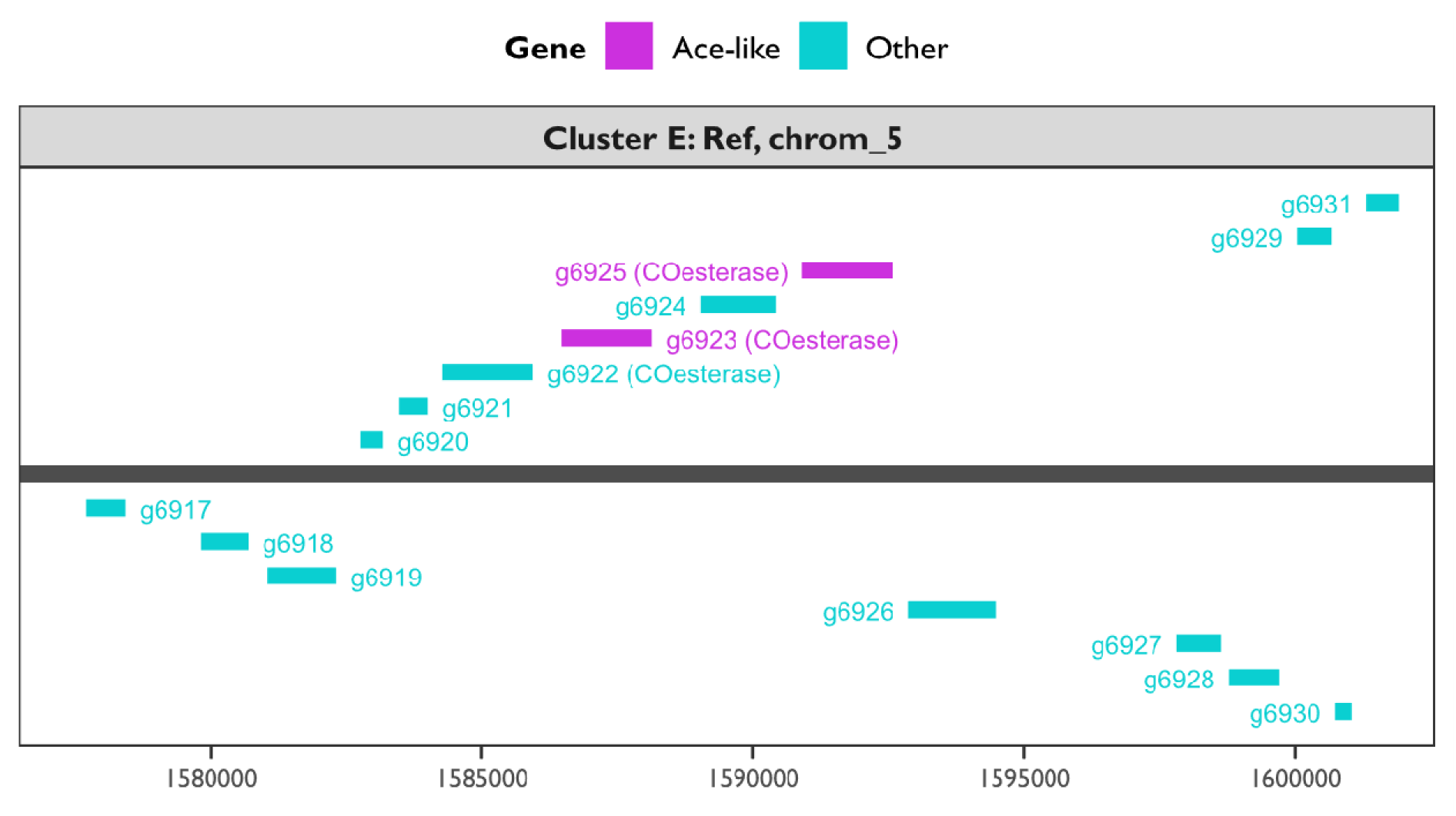
Genomic context around *ace*-like genes in cluster E (as per Figure 7). Data support 2 putative paralogous genes on the same genomic sequence. The *x*-axis represents the base coordinate on the assembled chromosome (or contig). The grey horizontal bar represents the genomic sequence. The magenta bars represent the *ace*-like genes. The turquoise bars represent other genes within 10,000 bp of the *ace*-like genes. Genes *above* the grey genomic sequence are those on the positive strand, whereas those *below* the grey genomic sequence are those on the negative strand. Panels separate different chromosomes (or contigs).

**Figure S11.**
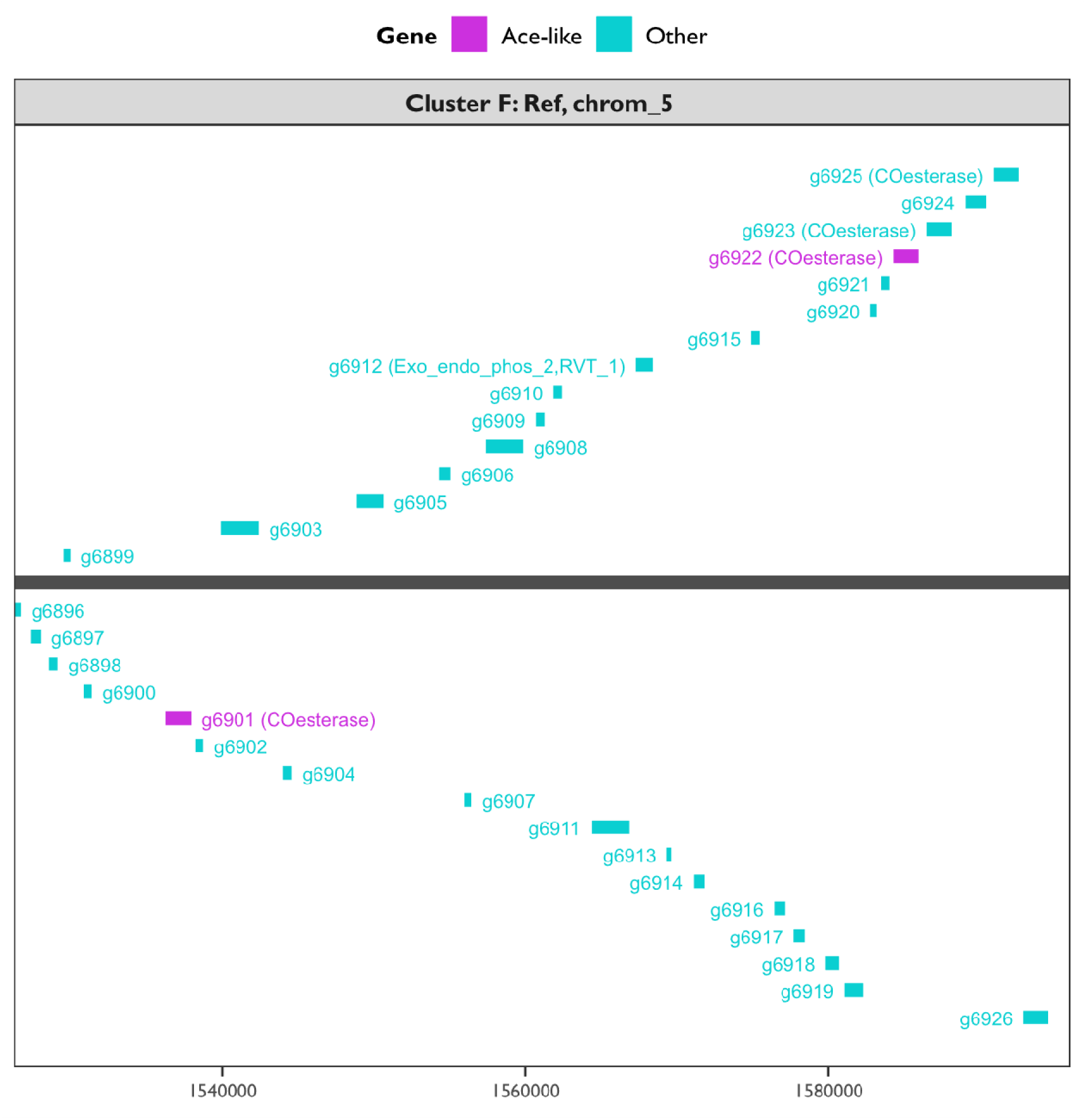
Genomic context around *ace*-like genes in cluster F (as per Figure 7). Data support two putative orthologous genes artificially split on different genomic sequences. The *x*-axis represents the base coordinate on the assembled chromosome (or contig). The grey horizontal bar represents the genomic sequence. The magenta bars represent the *ace*-like genes. The turquoise bars represent other genes within 10,000 bp of the *ace*-like genes. Genes *above* the grey genomic sequence are those on the positive strand, whereas those *below* the grey genomic sequence are those on the negative strand. Panels separate different chromosomes (or contigs).

**Figure S12.**
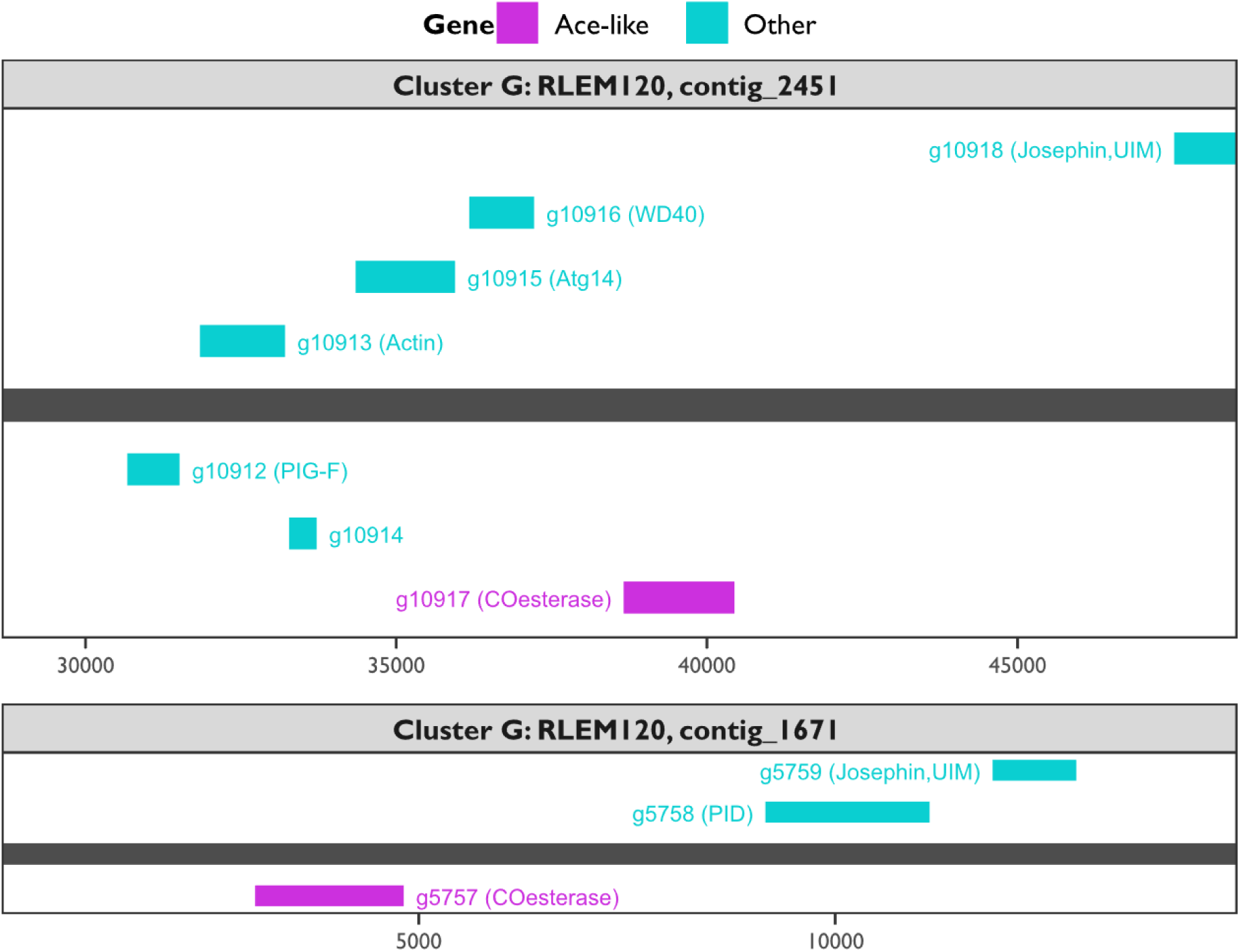
Genomic context around *ace*-like genes in cluster G (as per Figure 7). Data support 2 putative orthologous genes artificially split on different genomic sequences. The *x*-axis represents the base coordinate on the assembled chromosome (or contig). The grey horizontal bar represents the genomic sequence. The magenta bars represent the *ace*-like genes. The turquoise bars represent other genes within 10,000 bp of the *ace*-like genes. Genes *above* the grey genomic sequence are those on the positive strand, whereas those *below* the grey genomic sequence are those on the negative strand. Panels separate different chromosomes (or contigs).

**Figure S13.**
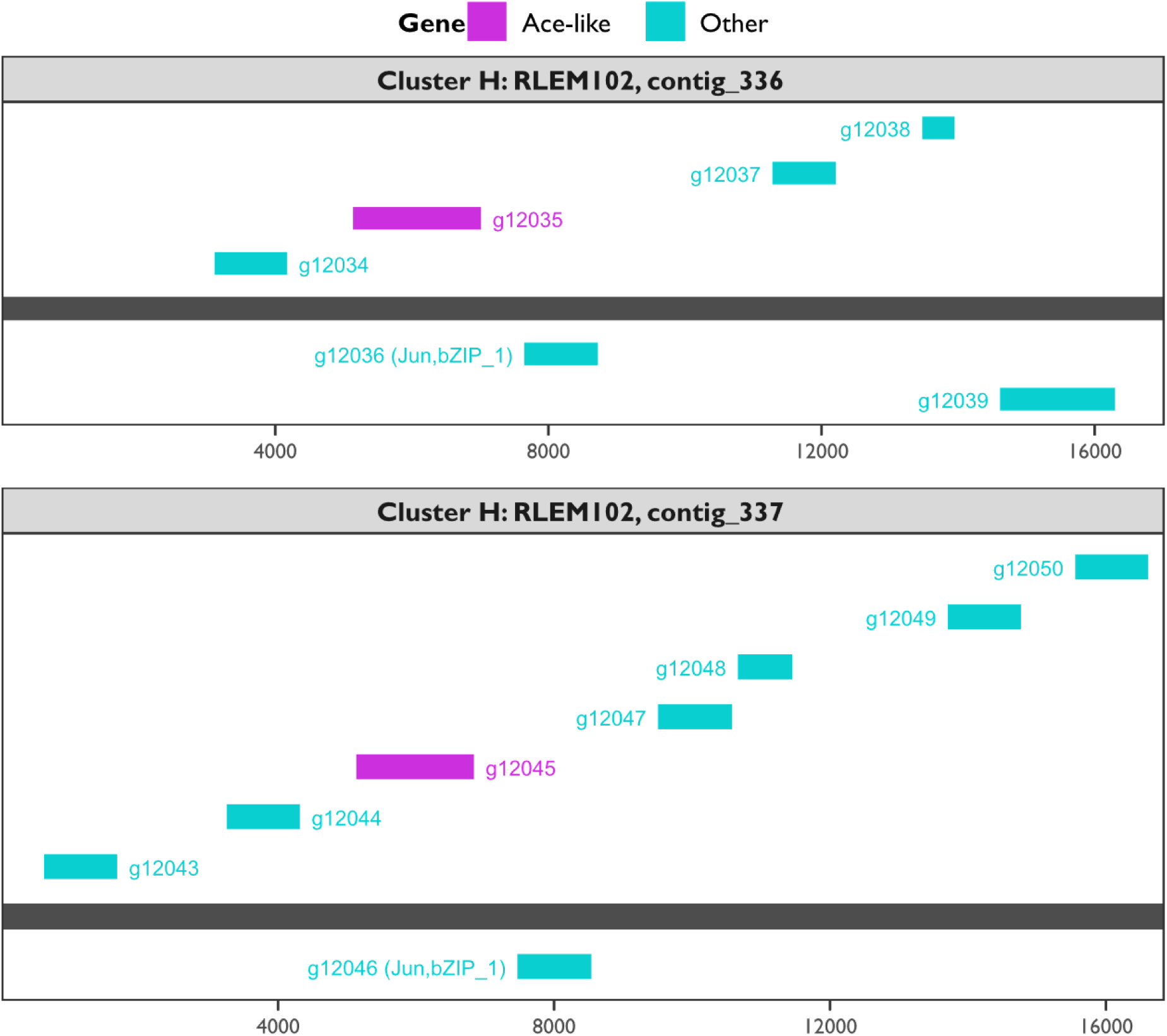
Genomic context around *ace*-like genes in cluster H (as per Figure 7). Data support 2 putative orthologous genes artificially split on different genomic sequences. The *x*-axis represents the base coordinate on the assembled chromosome (or contig). The grey horizontal bar represents the genomic sequence. The magenta bars represent the *ace*-like genes. The turquoise bars represent other genes within 10,000 bp of the *ace*-like genes. Genes *above* the grey genomic sequence are those on the positive strand, whereas those *below* the grey genomic sequence are those on the negative strand. Panels separate different chromosomes (or contigs).

**Table S1.**
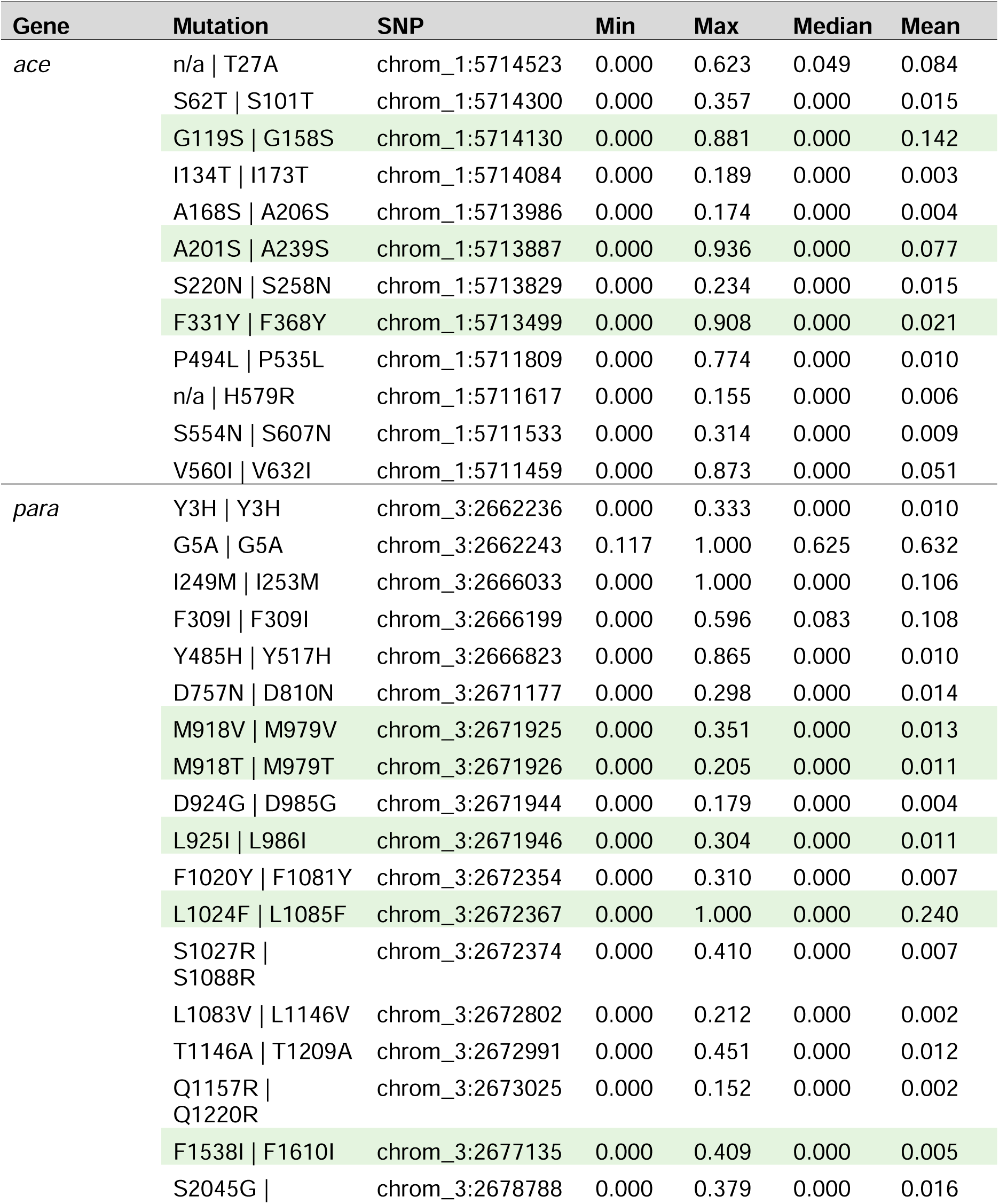
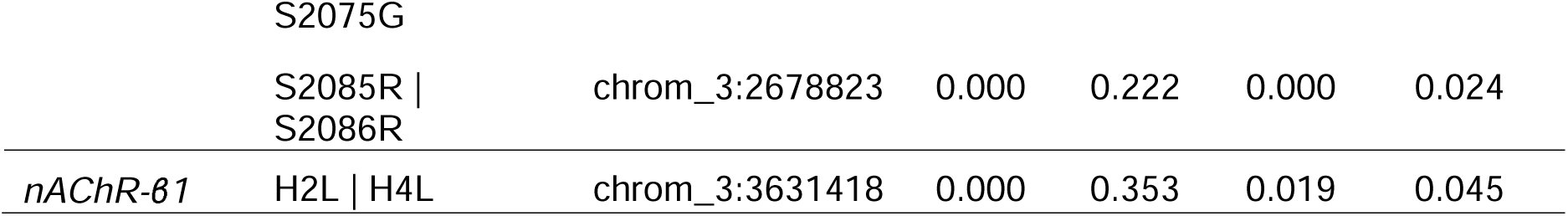
Non-synonymous mutations in pesticide target genes of *Halotydeus destructor*. Mutations are reported as canonical reference numbering first (*Torpedo californica* for *ace*, *Musca domestica* for *para*; and *Myzus persicae* for *nAChR*), *H. destructor* numbering second, separated with a bar. An ‘n/a’ indicates an equivalent position does not exist. SNP loci are reported as the chromosome and position. Highlighted rows indicate mutations that have been linked to target-site resistance in other arthropods. Mutation frequencies are reported as their min, max, median and mean across 190 populations.

**Table S2.** Analysis of deviance for models testing the association between resistance with *ace* coverage (read depth) and the frequency of the G119S and A201S mutations in *Halotydeus destructor*. *P*-values significant at a threshold of α = 0.05 are shaded pink.

| Chemical | State | Term | LR Chisq | Df | Pr(>Chisq) |
| --- | --- | --- | --- | --- | --- |
| Chlorpyrifos | WA | Read depth | 0.406 | 1 | 0.524 |
|  |  | G119S freq | 141.772 | 1 | <0.001 |
|  |  | A201S freq | 0.258 | 1 | 0.612 |
|  | SA | Read depth | 5.792 | 1 | 0.016 |
|  |  | G119S freq | 134.504 | 1 | <0.001 |
|  |  | A201S freq | 5.722 | 1 | 0.017 |
|  | VIC | Read depth | 0.024 | 1 | 0.878 |
|  |  | G119S freq | 0.134 | 1 | 0.715 |
|  |  | A201S freq | 0.171 | 1 | 0.679 |
| Omethoate | WA | Read depth | 71.819 | 1 | <0.001 |
|  |  | G119S freq | 3.626 | 1 | 0.057 |
|  |  | A201S freq | 4.296 | 1 | 0.038 |
|  | SA | Read depth | 0.068 | 1 | 0.794 |
|  |  | G119S freq | 22.768 | 1 | <0.001 |
|  |  | A201S freq | 16.093 | 1 | <0.001 |
|  | VIC | Read depth | 0.221 | 1 | 0.639 |
|  |  | G119S freq | 0.573 | 1 | 0.449 |
|  |  | A201S freq | 0.377 | 1 | 0.539 |

**Table S3.** Partial regression coefficients for models testing the association between resistance with *ace* coverage (read depth) and the frequency of the G119S and A201S mutations in *Halotydeus destructor*.

| Chemical | State | Coeff | Estimate | SE |
| --- | --- | --- | --- | --- |
| Chlorpyrifos | WA | Intercept | −2.58 | 0.23 |
|  |  | Read depth | −0.13 | 0.2 |
|  |  | G119S | 1.77 | 0.2 |
|  |  | A201S | −0.11 | 0.22 |
|  | SA | Intercept | −1.65 | 0.17 |
|  |  | Read depth | −0.37 | 0.15 |
|  |  | G119S | 1.99 | 0.2 |
|  |  | A201S | 0.36 | 0.15 |
|  | VIC | Intercept | −6.12 | 0.89 |
|  |  | Read depth | −0.18 | 1.17 |
|  |  | G119S | 1.15 | 2.87 |
|  |  | A201S | −1.3 | 3.36 |
| Omethoate | WA | Intercept | −3.58 | 0.33 |
|  |  | Read depth | 1.27 | 0.16 |
|  |  | G119S | 0.31 | 0.16 |
|  |  | A201S | 0.39 | 0.17 |
|  | SA | Intercept | −1.18 | 0.22 |
|  |  | Read depth | −0.06 | 0.23 |
|  |  | G119S | 1.2 | 0.26 |
|  |  | A201S | 0.73 | 0.19 |
|  | VIC | Intercept | −4.84 | 0.33 |
|  |  | Read depth | −0.22 | 0.47 |
|  |  | G119S | 0.87 | 1.15 |
|  |  | A201S | −0.7 | 1.16 |

